# The HES1-SOD1 Antagonism Shapes Senescence Heterogeneity and Impacts Metastatic Relapse of Circulating Tumor Cells

**DOI:** 10.64898/2026.08.19.745859

**Authors:** Guanyin Huang, Xiaolong Xu, Binyu Zhang, Mao Zhao, Yixin Cheng, Boxi Zhao, Shuqian Zheng, Xuefei Liu, Songyan Yu, Liuyang Wang, Jianyang Hu, Chunhao Long, Yihuizhi Zhang, Yuling Sheng, Siyuan Xia, Lin Zeng, Hao Yu, Hui Yang, Jialing Liu, Yi Lu, Jian Zhang, Weineng Feng, Meng Xu, Weinan Guo, Xin Hong

## Abstract

Circulating tumor cells (CTCs) encounter multiple challenges within the blood microenvironment, including oxidative stress, flow shear forces, and immune surveillance, often leading to anoikis. Recently, a transitional state of CTC senescence has been identified, contributing to metastatic inefficiency. However, the molecular mechanisms linking senescent CTCs to disease relapse remain to be defined. By integrating a genetic model of cortactin knockdown-induced CTC senescence with single-cell multi-omic analyses, we revealed two distinct senescent CTC subpopulations marked by HES1 expression levels. These HES1^low^ and HES1^high^ subpopulations exhibited differential evolutionary trajectory dynamics and unique molecular and metabolic signatures, which were significantly correlated with adverse clinical outcome across several patient cohorts. HES1^low^ senescent CTCs displayed enhanced mitochondrial fitness, oxidative phosphorylation, and ROS-detoxifying capabilities, resulting in more efficient tumor regrowth with a pro-inflammatory and thrombotic phenotype when compared to the HES1^high^ group. Mechanistically, HES1 directly bound to the *Sod1* promoter and repressed its expression, leading to redox imbalance and mitochondrial dysfunction that were linked to weakened tumor regrowth capacity. Both senescent CTC subpopulations were broadly resistant to cytotoxic and targeted therapies, yet they showed elevated dependency on anti-apoptosis programs that make them susceptible to dual blockade by SOD1 inhibitor and the anti-senolytic drug ABT737 *in vivo*. Finally, in a prospective cohort of on-treatment melanoma patients, HES1⁺ senescent CTCs were highly enriched in patients with progressive disease. Thus, the HES1-SOD1 antagonism shapes CTC senescence heterogeneity and contributes to differential tumor relapse, which can be therapeutically explored for eliminating residual metastatic disease.

**GRAPHIC ABSTRACT:** 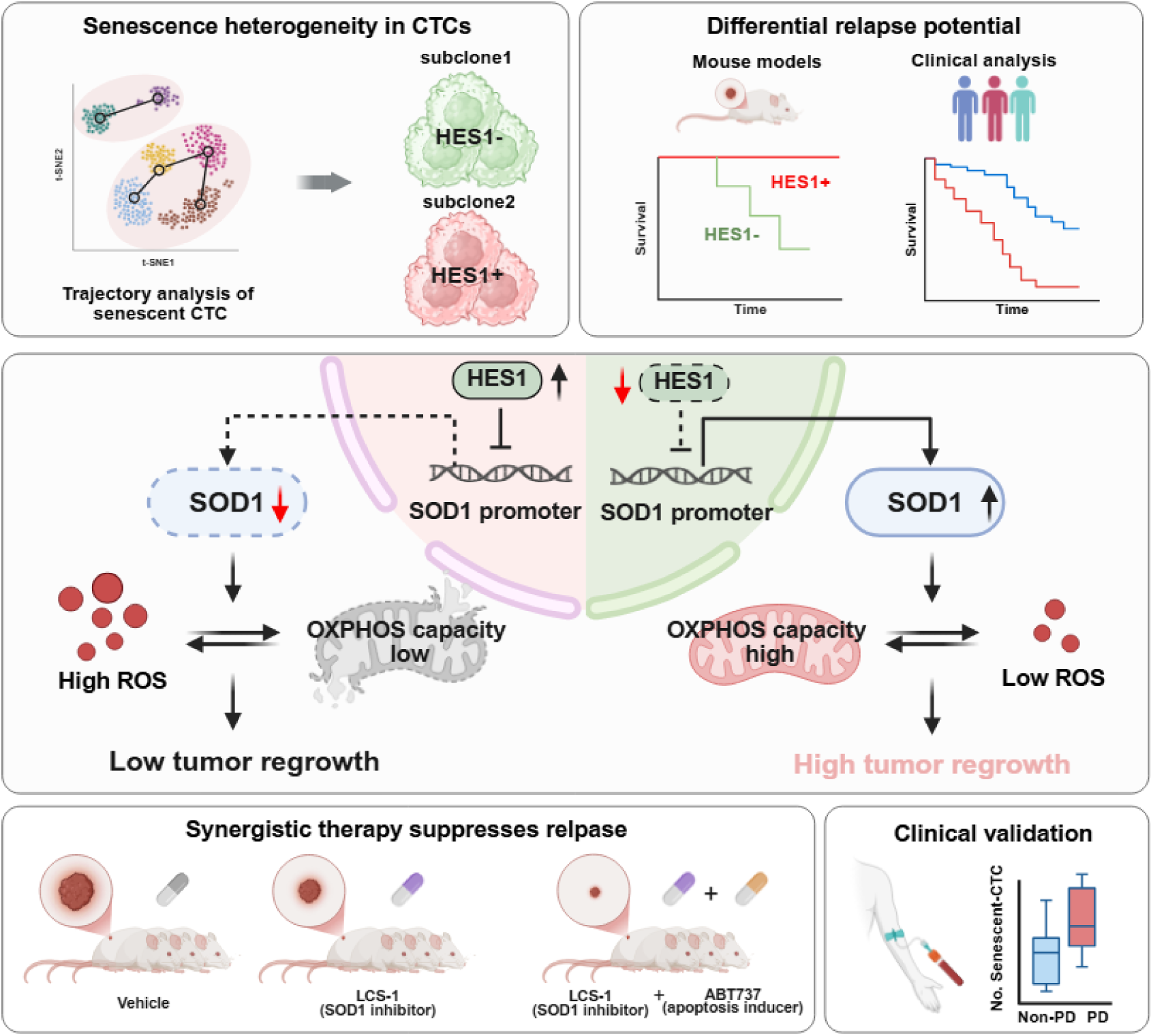

Trajectory analysis of senescent CTCs uncovers substantial heterogeneity, resolving two major subclones defined by HES1 expression (HES1⁺ and HES1⁻) with distinct molecular and metabolic programs. These subclones exhibit clinically prognostic correlation and linked to metastatic relapse. Mechanistically, HES1 and SOD1 are antagonistically regulated to control mitochondrial fitness and ROS homeostasis. HES1 represses the SOD1 expression, which elevates ROS and impairs OXPHOS capacity, thereby constraining tumor regrowth. Preclinical model showed the SOD1 inhibitor LCS-1 combined with the apoptosis inducer ABT737 synergistically suppresses senescent CTC-driven tumor relapse *in vivo*. Clinically, melanoma patients with progressive disease (PD) harbor significantly more senescent CTCs than non-PD patients, highlighting senescent CTCs as a noninvasive biomarker for therapeutic resistance and disease monitoring.

## INTRODUCTION

Circulating tumor cells (CTCs) are considered pre-metastatic precursors, yet only a minor fraction successfully colonizes distant organs, enter dormancy, and eventually give rise to lethal metastatic relapse^1^. A wide range of stimulus, including oxidative stress, flow shear forces, immune surveillance, and therapeutic interventions can drive cancer cells into a senescent state^2,3^. Although senescence is originally defined as a state of irreversible cell cycle arrest, cancer cells appear to hijack this mechanism to enable short-term dormancy and disease recurrence ^3–5^. Recent studies have demonstrated that senescent tumor cells can re-awaken and contribute to metastatic recurrence^6–8^. Whether and how pre-metastatic senescent CTCs engage in disease relapse remains an open question.

Heterogeneity is a hallmark of cancer and manifests at phenotypic and functional levels, encompassing features linked to tumor microenvironment compositions, cellular plasticity, genetic and epigenetic alterations, and therapeutic adaptations ^9–11^. Recent evidence has revealed that senescent cells, particularly senescent tumor cells, also exhibit pronounced heterogeneity^4,11,12^. Some reports suggest that senescent tumor cells can activate stemness programs and re-enter the cell cycle to drive relapse^6–8,13^, whereas others show that recurrence arises from a drug-tolerant persister (DTP) population rather than from bona-fide senescent cells^14^. These conflicting observations underscore an urgent need to decipher the senescence platicity of tumor cells and identify lethal subpopulations that contribute to disease progression. The molecular mechanisms governing CTC senescence heterogeneity and its impact on metastatic relapse remain to be addressed.

HES1 is a transcriptional repressor downstream of Notch signaling, originally discovered as a homologue of the Drosophila *hairy* and *Enhancer of split* genes^15,16^.

HES1 functions in a broad range of biological processes, including embryonic development^15,16^, cell fate determination^17^, stem cell maintenance^18^, and oncogenesis^19^. Notably, HES1 has been implicated as a key factor that safeguards quiescent fibroblast against irreversible cell cycle arrest and inappropriate differentiation^20–22^. Despite its established role in stemness regulation, whether HES1 participates in the senescence regulation in CTCs has not been reported.

Here, by integrating a genetic CTTN knockdown-induced CTC senescence model and multi-omic analyses, we identified HES1+ and HES1- senescent subpopulations exhibiting distinct molecular and mitochondrial metabolic features. Molecularly, HES1 directly repressed SOD1 transcription, and the antagonism between HES1 and SOD1 impacted redox balance and tumor regrowth potential. Combination treatment with LCS-1 and ABT737 delayed tumor recurrence both *in vitro* and *in vivo*. Finally, as a proof of principle, we demonstrated that HES1^+^ senescent CTCs were significantly enriched in a prospective cohort of progressing melanoma patients undergoing immunotherapy.

## RESULTS

### Single-cell transcriptomic landscape of senescent melanoma CTCs

We employed a genetically faithful CTTN knockdown (KD) system to model senescence in melanoma CTCs^2^. CTTN KD resulted in an increased proportion of β-galactosidase-positive (β-gal^+^) CTCs, inducing cell cycle arrest and triggering a senescent phenotype as described previously (Figure 1A-D). This was evidenced by the upregulation of classical senescence markers (p53 and p21) and downregulation of lamin B1 (LMNB1), collectively leading to cellular senescence (Figure 1E)^2^. To investigate the molecular heterogeneity of senescent CTC subclones, we further performed single-cell transcriptomic profiling on wild type (Control) and CTTN-KD

**Figure 1.**
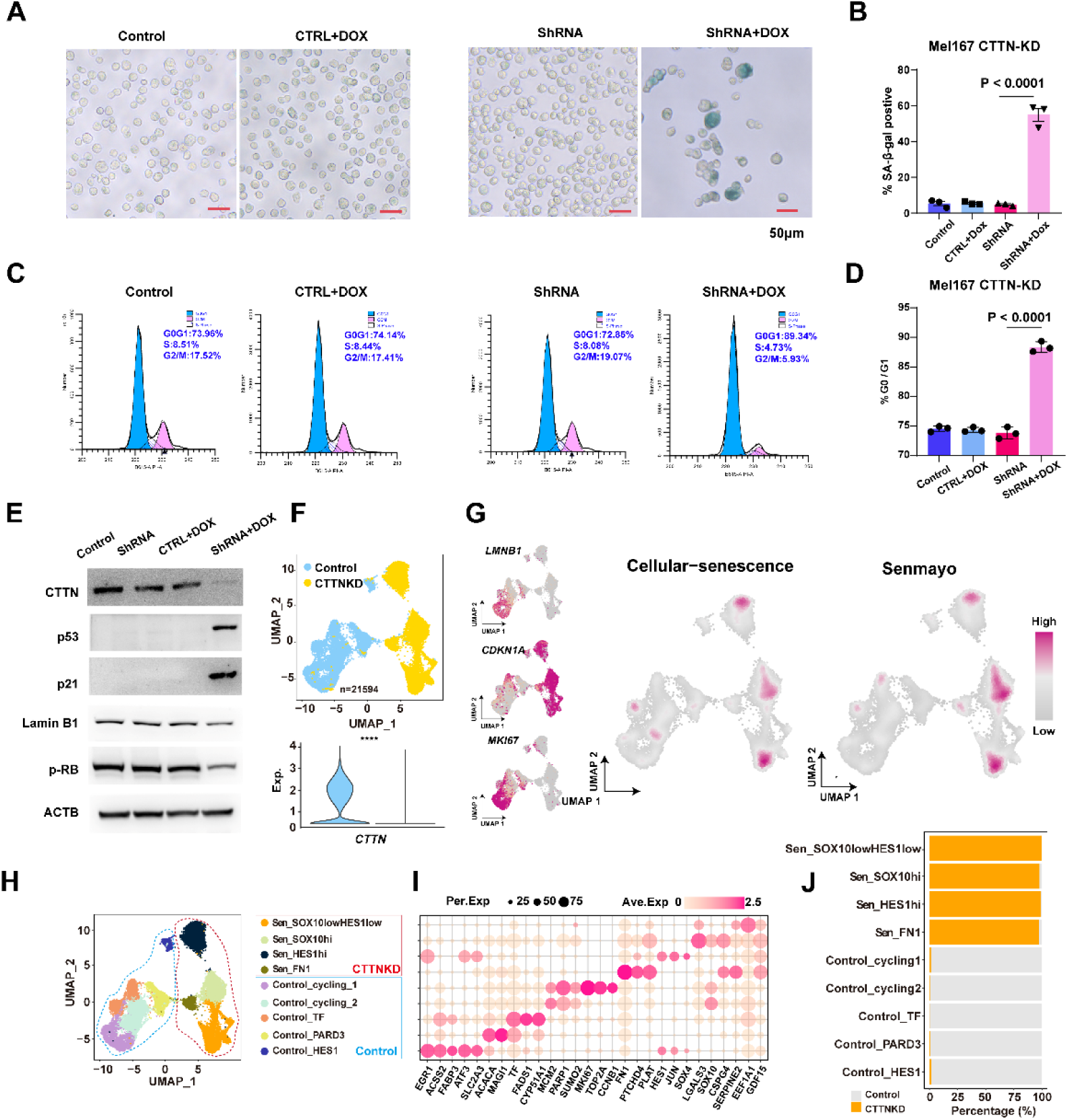
Single-cell transcriptomic landscape of senescent melanoma CTCs. (A) Representative image shows the SA-β-gal staining of Mel-167 cells with or without CTTN-KD for 6 days. (B) Quantification of the percentage of SA-β-gal–positive cells. *n* =3 independent experiments. (C) Cell-cycle analysis of Mel-167 CTCs with or without CTTN KD. (D) Quantification of G0–G1-, S-, and G2–Mphase of cells in D. *n* = 3 independent experiments. (E) Western blotting shows protein levels of cortactin, p53, p21, pRB, and Lamin B1 proteins in Mel-167 cells. ACTB as loading control. (F) UMAP visualization of wild-type and CTTN-KD melanoma CTCs (following call senescent melanoma CTCs) (top). Yellow highlights senescent melanoma CTCs while Blue represents wild-type melanoma CTCs. Violin plot shows the mRNA expression of CTTN in each group (bottom). (G) UMAP shows the mRNA expression level of classical senescent associated markers (LMNB1, CDKN1A, MKI67) and pathways (Cellular senescence and Senmayo) among wild-type and senescent CTCs subpopulations. (H) UMAP visualization of subpopulations of wild-type and senescent melanoma CTCs. Each dot indicates a single cell. Color-coded for clusters. (I) Dot plot shows genes highly expressed in each subpopulation presented in Figure 1H. (J) Bar plot shows relative proportions of wild-type and senescent melanoma CTCs in each subpopulation. **B, D,** and **F,** Data are represented as mean ± SEM. **B,** and **D,** Statistical significance was calculated by two-way ANOVA with the Dunn *post hoc* test; **F,** Statistical significance was calculated by Wilcoxon test. ****, *P* < 0.0001. Dox, doxycycline.

CTCs. After quality control, a total of 21594 cells passed the quality control (Figure 1F; Figure S1A). Our analysis revealed the expressing distribution of classical senescence markers, including the elevation of p53 and p21 and downregulation of lamin B1 (LMNB1) and MKI67 at single-cell resolution. CTTN KD cells also showed strong enrichment of senescence-associated pathways, including “Cell-cycle arrest”, “Senmayo”, and “Cellular-senescence” when compared to the control group (Figure 1G; Figure S1B-D). Furthermore, CTTN-KD CTCs downregulated lipid metabolism-related genes such as *ACLY*, *ACSS2*, and *FASN* and pathways involved in cholesterol homeostasis (Figure 1G; Figure S1E-F). Unsupervised clustering analyses were performed on WT and CTTN-KD CTCs, revealing 9 subpopulations with distinct transcriptional profiles, including 5 subpopulations within the WT non-senescent CTCs and 4 senescent subpopulations in CTTN-KD CTCs (Figure 1H-J). These data demonstrated that CTTN depletion resulted in transcriptionally heterogeneous senescent CTC subpopulations with distinct molecular profiles.

### Evolutionary trajectory analysis uncovers two distinct senescent CTC subpopulations associated with poor clinical outcome

To investigate the molecular heterogeneity and evolutionary dynamics of distinct senescent CTC subgroups, trajectory analysis was performed. The results showed that there were likely two transition routines for CTCs to enter senescent states (Figure 2A-B; Figure S2A-B). The subpopulation transition in Routine1 was Control->Sen_FN1->Sen_SOX10^high^->Sen_SOX10^low^HES1^low^ (Sen_HES1^low^ in short). The Routine 2 transition was from Control_HES1 to Sen_HES1^high^. Thus, Sen_HES1^low^ and Sen_HES1^high^ were likely two independently evolved terminal senescent subgroups (Figure 2A-B; Figure S2A-B). We further generated two distinct molecular signatures derived from these two senescent CTC subpopulations as “Sen_HES1low signature” and “Sen_HES1high signature”, and explored their prognostic values in monitoring melanoma disease progression. In an independent melanoma cohort (GSE15605)^23^, both senescence-associated CTC signature scores were significant higher in metastasis samples compared to primary tumor or normal tissue samples (Figure S2C). In another two independent patient cohorts (GSE19234^24^ and GSE65904^25^), Kaplan-Meier survival analysis showed that patients with higher signature scores exhibited significantly shorter distant metastasis-free survival (DMFS) than those with low signature scores in both categories (Figure 2C). Furthermore, in TCGA-SKCM cohort, patients with higher signature scores also showed significantly shorter disease-specific survival (DSS) than those with low signature scores (Figure S2D). These clinical analyses from independent cohorts implicated that both Sen_HES1^low^ and Sen_HES1^high^ CTC subpopulations were clinically prognostic and may be functionally linked to metastatic relapse (Figure 2C; Figure S2C-D).

**Figure 2.**
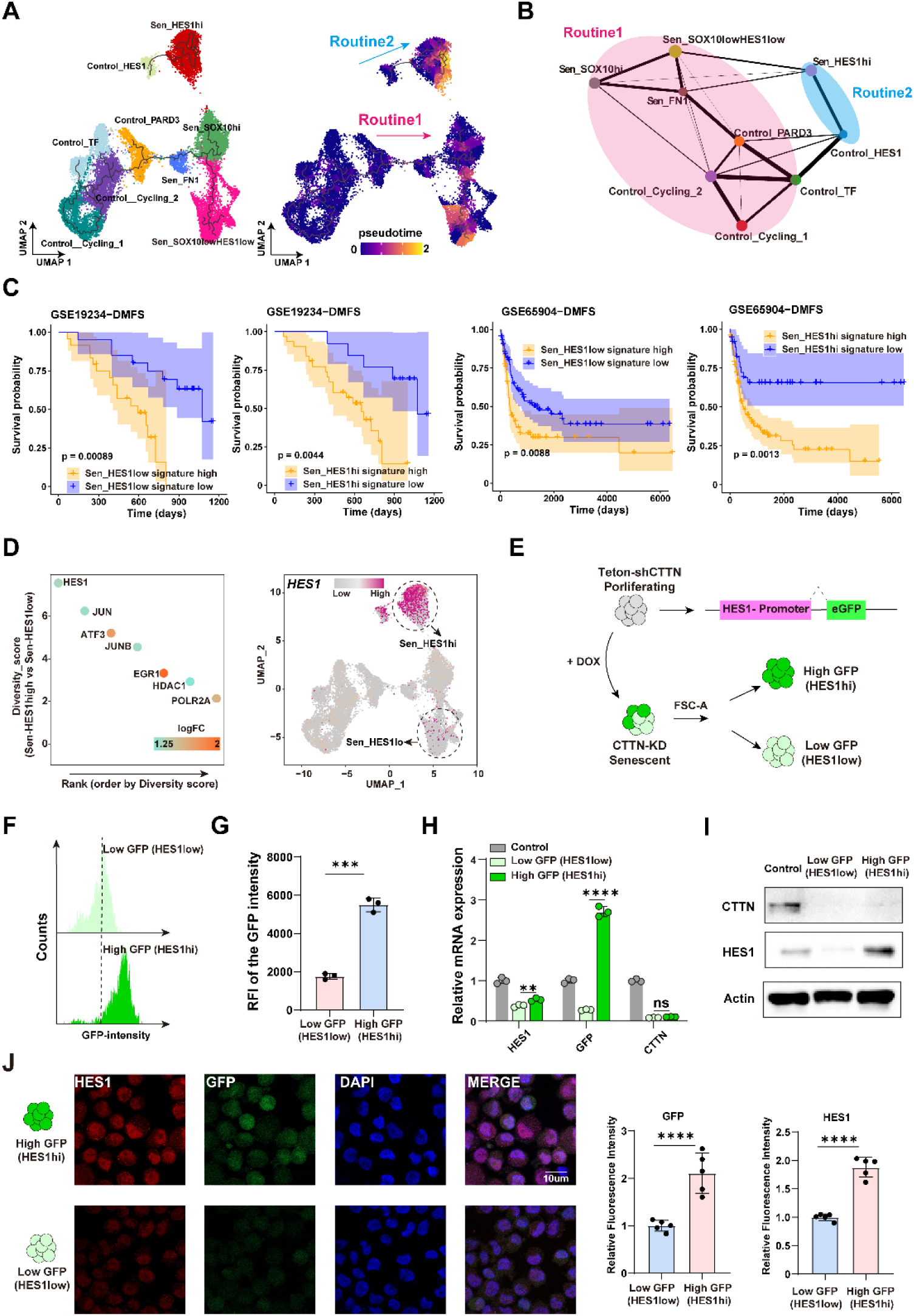
Evolutionary trajectory analysis uncovers two distinct senescent CTC subpopulations associated with poor clinical outcome. (A) Trajectory analysis of wild-type and senescent melanoma CTCs was performed by Monocle 3, colored by inferred pseudotime. (B) Partition-based graph abstraction (PAGA) analysis of 9 melanoma CTCs subtypes. (C) Distant metastasis free survival of melanoma patients in the GSE19234 (left) and GSE65904 (right) cohort based on Sen_HES1^low^ and Sen_HES1^high^ subpopulations signatures expression. (D) Regulator factors highly expressed in Sen_HES1^high^ subpopulations (left). *Y*-axis indicates the diversity distribution within Sen_HES1^high^ subpopulations; Score was calculated as the ratio of gene expression positive cells in Sen_HES1^high^ subpopulations versus out of Sen_HES1^high^ subpopulations (Sen-HES1^high^ versus Sen-HES1^low^), genes ordered by diversity distribution along the *X*-axis. UMAP shows the mRNA expression level of *HES1* among wild-type and senescent CTCs subpopulations (right). (E) Schematic diagram of HES1-promoter-eGFP report system to isolate the Sen_HES1^high^ and Sen-HES1^low^ subpopulations following CTTN-KD in CTCs. (F) Representative flow cytometry plots show GFP intensity-based gating of HES1^low^ and HES1^high^ senescent CTCs. (G) Bar graph shows the relative fluorescence intensity (RFI) of GFP shown in Figure 2F. *n* = 3 independent experiments. (H) Relative mRNA expression of *HES1*, *GFP*, *CTTN* determined by RT-qPCR in HES1^low^, HES1^high^ senescent CTCs and wild-type CTCs. *n* = 3 independent experiments. (I) Western blotting shows protein levels of CTTN and HES1 in HES1^low^, HES1^high^ senescent CTCs and wild-type CTCs. Actin as loading control. (J) Multiplexed immunofluorescence (mIF) staining of HES1^low^, HES1^high^ senescent CTCs (left) showing HES1 (red), GFP (green), DAPI (blue). Quantification of relative fluorescence intensity of HES1 and GFP in HES1^low^, HES1^high^ senescent CTCs (right). *n* = 5 independent experiments. Scale bar, 10 μm. **G, H,** and **J,** Data are represented as mean ± SEM. **G, H,** and **J,** Statistical significance was calculated by a two-sided Student *t* test; **C,** statistical significance was calculated by log-rank test. NS, not significant; **, *P* < 0.01; ***, *P* < 0.001; ****, *P* < 0.0001. Dox, doxycycline.

To identify key regulatory factors (RF) including transcription factors and epigenetic regulators that are highly enriched in these two terminal senescent CTC subpopulations, we conducted a four-way differential RF analyses between Routine1 and Routine2 in control CTCs versus the two distinct senescent CTC subpopulations (Figure S2E). Interestingly, CTCs in routine2 overexpressed RFs such as *EGR1*, *HES1*, *KLF4* related to early stress response and stemness regulations (Figure S2E). In contrast, CTCs in routine1 upregulated RFs such as *HIF1A*, *CTNNB1*, *PRKDC*, and pathways involved in hypoxia stress and WNT pathway (Figure S2E). Among them, we focused on HES1 because it was the most differentially expressed RF between Sen-HES1^high^ and Sen-HES1^low^ subgroup (Figure 2D).

To experimentally assess the molecular functions of these two distinct senescent CTC subgroups, we constructed a HES1-promoter-eGFP reporter CTC line to isolate the Sen_HES1^high^ and Sen-HES1^low^ CTC subpopulations following CTTN KD in Mel-167 CTCs (Figure 2E-G). The HES1 promoter GFP expression intensity correlated well with endogenous HES1 expression level in isolated CTC subpopulations as measured by flow cytometry-based cell sorting (FACS) followed by HES1 mRNA and protein quantifications (Figure 2E-I). FACS-purifed GFP^high^ senescent CTCs showed substantially elevated *HES1* expression compared to GFP^low^ senescent CTCs (Figure 2H-I). Multiplexed immunofluorescence further confirmed that GFP^high^ senescent CTCs displayed higher protein expression of HES1 and GFP compared to the GFP^low^ group (Figure 2J). In addition, we verified the specificity of HES1 GFP reporter using a conventional control eGFP as a negative control. In comparison, we observed no significant differences in HES1 expression using Mel-167 CTCs expressing a control GFP reporter, indicating that the HES1 promoter GFP reporter expression is specifically responding to HES1 transcriptional activities in the GFP reporter CTC line (Figure S2F-G). Together, we have established a sensitive HES1 promoter GFP reporter system to effectively label and isolate Sen_HES1^high^ and Sen-HES1^low^ CTCs.

### HES1^high^ and HES1^low^ senescent CTC subpopulations exhibit differential regrowth potential

To test the functional properties of these two senescent CTC subpopulations, we first examined proliferation and colony-forming capabilities *in vitro*. The HES1^low^ senescent CTCs exhibited significantly enhanced 2-D proliferation and 3-D colony formation capabilities than the HES1^high^ group (Figure 3A-B). To determine the tumorigenic potential of senescent CTC subpopulations *in vivo*, we subcutaneously implanted HES1^low^ and HES1^high^ senescent CTCs into immunocompromised NCG mice and monitored tumor growth over time. Remarkably, both senescent subpopulations initiated tumor regrowth after 5-weeks of implantation, and Sen_HES1^low^ exhibited significantly higher tumor growth rate than the Sen_HES1^high^ CTCs, yielding much larger tumors with nearly twice the mass of their Sen_HES1^high^ counterparts (Figure 3C-E; Figure S3A). To assess CTC-mediated metastatic potential, we injected luciferase-labelled senescent CTC subpopulations into the tail vein and monitored lung colonization by *in vivo* bioluminescence (BLI) imaging (Figure 3F). The results showed revealed visible lung metastasis in mice in the Sen_HES1^low^ group (3 of 5, 60%), compared with the Sen_HES1^high^ group (1 of 5, 20%, p = 0.07, Figure 3F). We observed a trend of reduced mice survival in the mice group bearing Sen_HES1^low^ tumors when compared to the Sen_HES1^high^ tumor-bearing mice (p=0.08, Figure 3G). While the Sen_HES1^low^ tumor-bearing mice start to die at around 60 days post tail-vein injection, the Sen_HES1^high^ tumor-bearing mice survived until the experimental endpoint (∼160 days) (Figure 3G). H&E staining revealed that lung tissues in decreased Sen_HES1^low^ tumor-bearing mice (3/3,100%) exhibited strong vascular thrombosis with large occlusive clots (>20 μm), whereas no significant vascular occlusion events were observed in the Sen_HES1^high^ tumor-bearing mice (Figure 3H-I; Figure S3B). Consistent with this observation, Sen_HES1^low^ subpopulations upregulated thrombosis- and inflammation-related factors such as *FN1*, *PLAT* and *ITGB3* and pathways related to inflammatory and coagulation pathways (Figure S3C-E). These results suggested that the Sen_HES1^low^ tumors may drive tissue inflammation and exhibited pro-thrombotic phenotypes during metastatic regrowth. Thus, these two senescent CTC subpopulations exhibited differential tumor regrowth and metastatic relapse potential.

**Figure 3.**
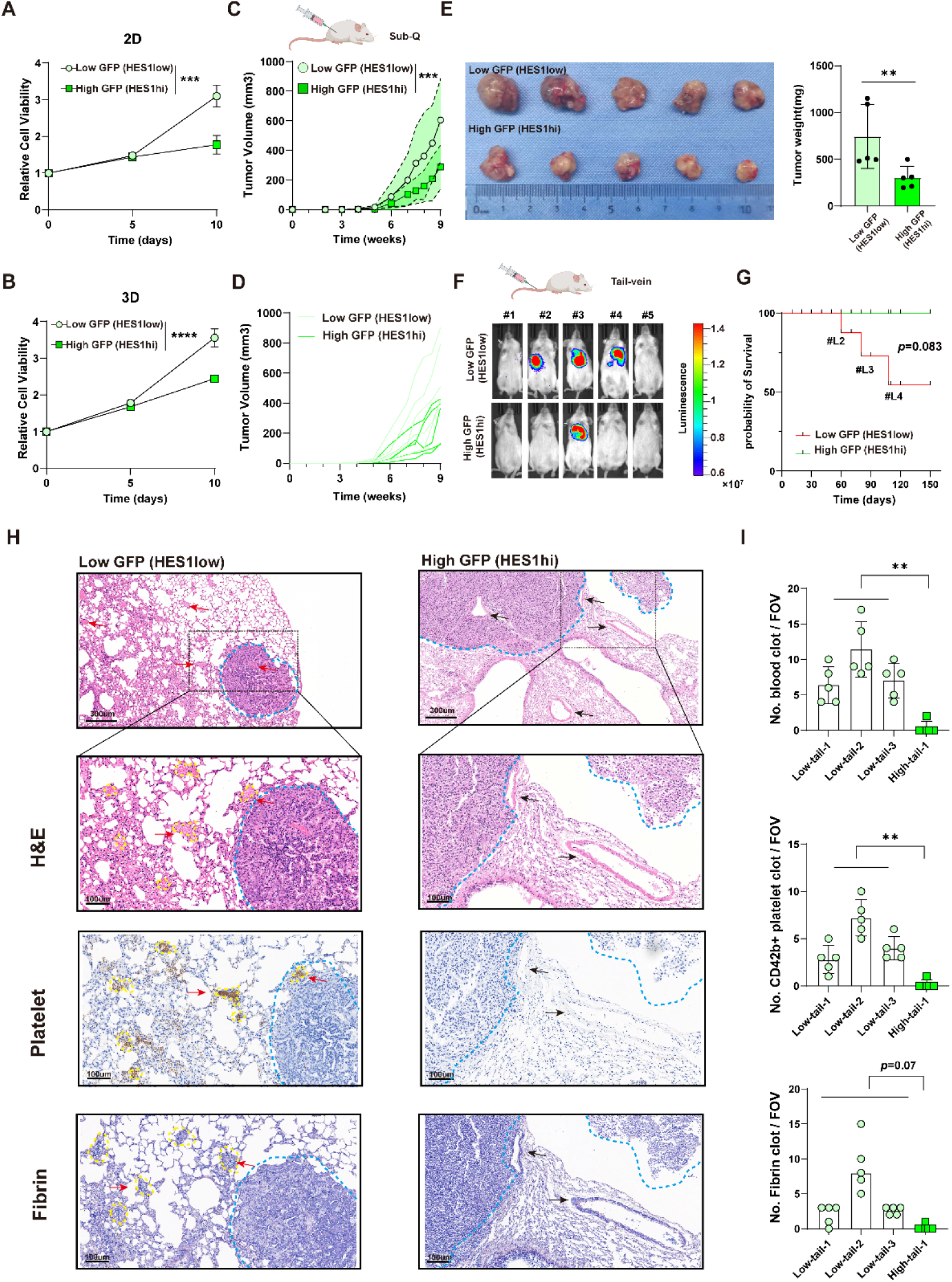
HES1^high^ and HES1^low^ senescent CTC subpopulations exhibit differential regrowth potential. (A) CCK8 proliferation assay of HES1^low^ and HES1^high^ senescent CTCs. (B) Colony formation rate of HES1^low^ and HES1^high^ senescent CTCs. (C-D) Tumor growth curve of HES1^low^ and HES1^high^ senescent CTCs subcutaneously injected into NCG mice. *N* =5 for each group. Each line represents individual mice in each group (D). (E) Tumor images (left) and paired tumor weight quantification (right) of the experiment shown in Figure 3C. (F) Tumor metastasis experiment established by tail vein injection of HES1^low^ and HES1^high^ senescent CTCs. The picture shows the In Vivo Imaging System (IVIS) images captured at the final time point before each mouse reached death or at the pre-defined experimental endpoint. *n*= 5 mice per group. (G) Survival curves of mice bearing HES1^low^ and HES1^high^ senescent CTCs in Figure 3F. (H) H&E staining (top), IHC staining (middle), and PATH staining (bottom) of lung metastasis of HES1^low^, HES1^high^ groups in Figure 3F. Lung metastasis were circuit with blue dash line. Red arrow represents vascular vessels with thrombosis. Black arrow represents vascular vessels without thrombosis. PATH stained for Fibrin. IHC stained for platelets (GPIb, CD42b). (I) Quantification of blood clot (top), CD42b^+^ platelet clot (middle), and fibrin clot (bottom) per FOV in lung tissues of HES1^low^, HES1^high^ groups in Figure 3H. *n*= 5 FOV per mouse. **A, B, C, E,** and **I,** Data are represented as mean ± SEM. **A, B, C, E,** and **I,** Statistical significance was calculated by a two-sided Student *t* test. **G,** statistical significance was calculated by log-rank test. **, *P* < 0.01; ***, *P* < 0.001; ****, *P* < 0.0001. FOV, field of view; PATH, phosphotungstic acid-haematoxylin; IHC, immunohistochemistry.

### Distinct mitochondrial metabolic capabilities are linked to the differential tumor regrowth potential of senescent CTC subpopulations

Given the differential tumor regrowth potential of Sen_HES1^low^ and Sen_HES1^high^ CTCs *in vitro* and *in vivo*, we further assessed gene expression and pathway enrichment in these two groups. Notably, we found that mitochondrial-coded genes and pathways associated with the mitochondrial functions were significantly enriched in the Sen-HES1^high^ subpopulations, which exhibits slower relapse (Figure S4A-B). This observation was further supported by elevated mitochondrial DNA content in the Sen_HES1^high^ CTCs, indicating increased total mitochondrial content (mitochondrial copy number) when compared to Sen_HES1^low^ CTCs (Figure S4C). Interestingly, our bioinformatic analyses further revealed that Sen_HES1^low^ subpopulation overexpressed genes such as *PKM*, *MDH2*, *ACLY* and metabolic pathways related to oxidative phosphorylation, TCA cycle, and fatty acid degradation (Figure 4A; Figure S4D-E). In contrast, Sen_HES1^high^ CTCs were enriched with metabolic pathways related to glycosaminoglycan biosynthesis, folate biosynthesis and one carbon pool by folate (1-C metabolism) (Figure 4A). Thus, it appeared Sen_HES1^high^ CTCs contained more mitochondria but exhibited weaker oxidative phosphorylation capabilities.

**Figure 4.**
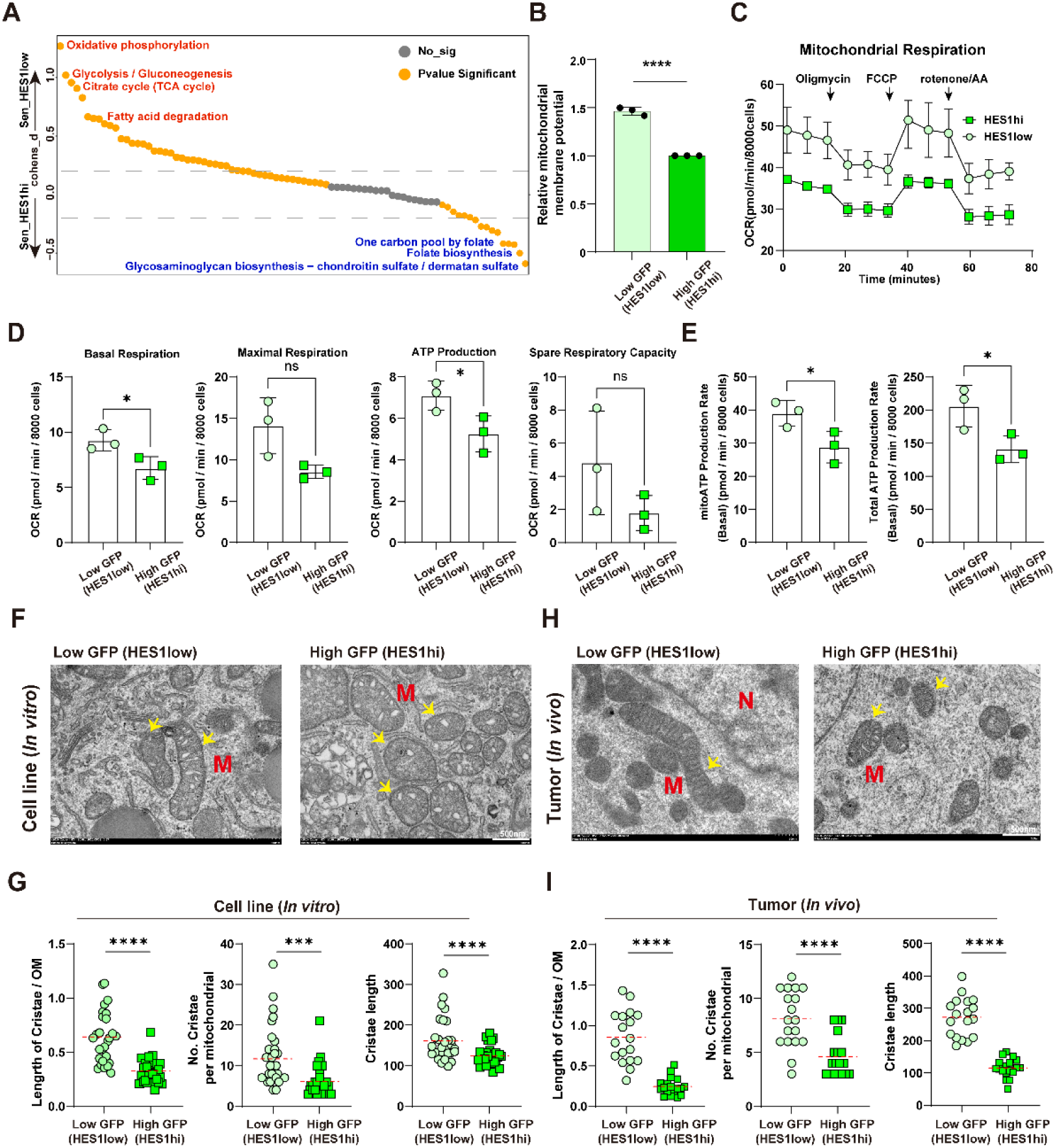
Distinct mitochondrial metabolic capabilities are linked to the differential tumor regrowth potential of senescent CTC subpopulations. (A) Metabolic pathway difference in Sen_HES1^low^ and Sen_HES1^high^ senescent CTCs. *Y*-axis indicates the cohens_d score diversity in metabolic pathway; Metabolic pathway ordered by cohens_d score along the *X*-axis. (B) Bar graph shows the relative mitochondrial membrane potential of HES1^low^, HES1^high^ senescent CTCs. *n* = 3 independent experiments. (C) Measurement of oxygen consumption rate (OCR) using Seahorse extracellular flux analyzer. (D) Calculations of basal respiration, ATP-linked respiration, maximal respiration, and spare respiratory capacity rates based on OCR data. (E) Mitochondrial ATP production rate and total ATP production rate were measured by Seahorse extracellular flux analyzer. (F) Representative images of mitochondrial morphology analysis by TEM in HES1^low^ and HES1^high^ senescent CTCs. (G) Quantification of length ratio of cristae to outer mitochondrial membrane, number of cristae per mitochondrion, and cristae length of HES1^low^, HES1^high^ senescent CTCs. (H) Mitochondrial morphology analysis by TEM in HES1^low^ and HES1^high^ relapsed tumors. (I) Quantification of length ratio of cristae to outer mitochondrial membrane, number of cristae per mitochondrion, and cristae length of HES1^low^, HES1^high^ relapsed tumors. **B, D, E, G,** and **I,** Data are represented as mean ± SEM. **B, D, E, G,** and **I,** Statistical significance was calculated by a two-sided Student *t* test; NS, not significant; *, *P* < 0.05; ***, *P* < 0.001; ****, *P* < 0.0001.

This apparent paradox prompted us to further examine the mitochondrial quantity and function in these subpopulations. JC-1 assay results demonstrated that Sen_HES1^low^ CTCs maintained significantly higher mitochondrial membrane potential, indicating greater oxidative capacity per mitochondrion, despite having fewer number of mitochondria. (Figure 4B; Figure S4G-H). To assess mitochondrial respiratory chain activity, we employed the seahorse assay to measure the oxygen consumption rate in the two senescent CTC subpopulations. Sen_HES1^low^ CTCs exhibited significantly elevated basal respiration, ATP-linked respiration, maximal respiration, and spare respiratory capacity when compared to the Sen_HES1^high^ group, indicating stronger mitochondrial oxidative metabolism (Figure 4D). Consistently, there were significantly higher basal oxidative phosphorylation and increased mitochondrial ATP production rate in Sen_HES1^low^ CTCs when compared to the Sen_HES1^high^ group (Figure 4E). These results indicated that despite there were fewer mitochondria in Sen_HES1^low^ CTCs, they exhibited more efficient oxidative metabolism and energy production than the Sen_HES1^high^ group.

One plausible explanation for the observed phenotype would be that the Sen_HES1^high^ CTCs retained more defective mitochondria than the Sen_HES1^low^ CTCs. To test this, we employed transmission electron microscopy (TEM) to capture ultrastructural changes of mitochondria in these two senescent subpopulations *in vitro* and *in vivo* (Figure 4F-I). Interestingly, there were significantly more aberrant mitochondrial structures, in the Sen_HES1^high^ CTCs than the Sen_HES1^low^ CTCs cultured *in vitro*, including rounded mitochondrial shape, swollen cristae morphology, and shortened cristae length (Figure 4F-G). Similar differences in mitochondrial morphologies and structures were observed in mice tumors derived from Sen_HES1^high^ (low tumor burden) and Sen_HES1^low^ CTCs (high tumor burden) (Figure 4H-I). These data suggested mitochondrial structural and functional heterogeneities in these two senescent CTC subpopulations may be linked differential tumor relapse potential.

Hence, we further explored the mitochondrial turnover and re-cycling system, the mitophagy pathway ^26^, in these CTC subpopulations. Interestingly, the Sen-HES1^low^ subpopulation exhibited significantly higher mitophagy activity when compared to Sen-HES1^high^ group, as assessed by the degree of colocalization between LAMP1 and TOM20 (Figure S4G-H). This was consistent with the observation that Sen-HES1^low^ senescent CTCs exhibited better mitochondria fitness and function than the Sen-HES1^high^ group (Figure 4; Figure S4A-4F).

Together, these results suggested that the mitochondrial function and homeostatic control, but not the total amount of mitochondria, was critical for enhanced regrowth potential of senescent CTCs.

### The HES1-SOD1 antagonism regulates ROS detoxification and regrowth potential of senescent CTCs

Given that redox status can directly reflect mitochondrial functionality ^2,27–29^, we next assessed ROS levels in these two senescent CTC subgroups. MitoSOX Red staining revealed a pronounced enrichment of mitochondrial superoxide in Sen_HES1^high^ CTCs compared to the Sen_HES1^low^ group (Figure 5A-B). Consistently, quantification of ROS levels in mice tumor tissues further confirmed that ROS accumulation was substantially higher in Sen_HES1^high^ CTC-derived tumors when compared to the Sen_HES1^low^ group (Figure 5C-D). Interestingly, single-sample GSEA (ssGSEA) revealed that Sen_HES1^low^ CTCs significantly upregulated the ROS detoxification pathway (Figure S5A). These data indicated that the two senescent CTC subpopulations exhibited differential degrees of ROS accumulation and clearance.

**Figure 5.**
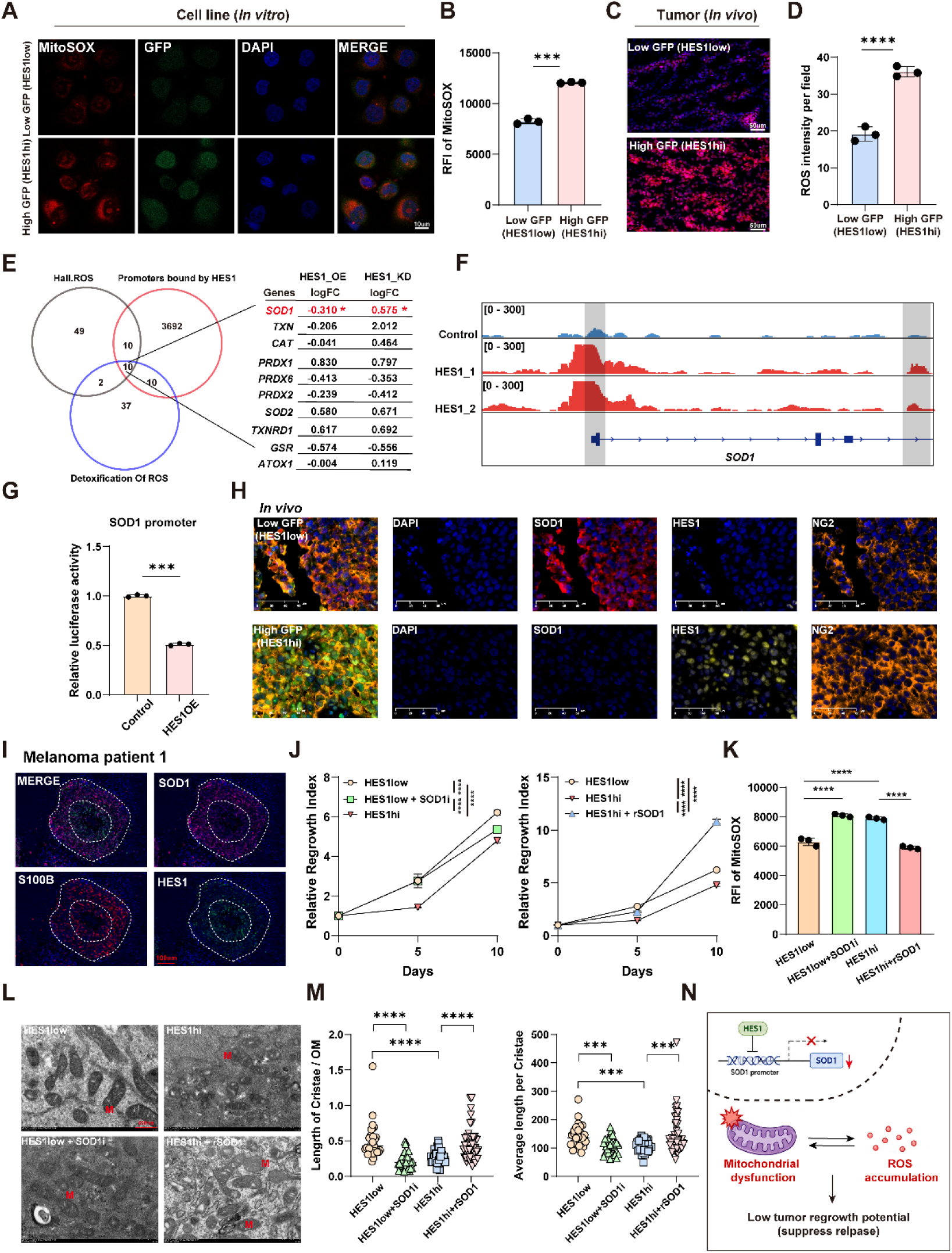
The HES1-SOD1 antagonism regulates ROS detoxification and regrowth potential of senescent CTCs. (A) Representative confocal images show MitoSOX staining (red), GFP expression (green), and DAPI nuclear staining (blue) in HES1-GFP^low and HES1-GFP^high senescent CTCs in vitro. Scale bar, 10 μm. (B) Quantification of RFI of MitoSOX in HES1^low^, HES1^high^ senescent CTCs. *n* = 3 independent experiments. (C) Represent confocal images of ROS (red) and DAPI (blue) staining in HES1^low^ and HES1^high^ relapsed tumors. Scale bar, 50 μm. (D) Quantification of ROS intensity per field in Figure 5C. *n* = 3 independent experiments. (E) Venn diagram (left) shows the overlap of genes among ROS pathway from Hallmark50, Detoxification of ROS pathway from Reactome, and the downstream targets of HES1 in MEL-167. Table (right) shows the relative mRNA expression level of 10 overlap genes in HES1-OE and HES1-KD versus wild-type MEL-167. (F) Integrative Genomics Viewer screenshot of HES1 binding peak on SOD1 in MEL-167. (G) Quantification analysis of the effect of *HES1* OE on the regulation of SOD1 promoter. (H) Representative images of mIHC shows DAPI (blue), SOD1 (red), HES1 (yellow), and NG2 (orange) in HES1^low^ and HES1^high^ tumors in Figure 3E. Each stain is shown separately and merged. Magnification, X40. (I) Representative images of mIHC shows SOD1 (pink), S100B (red), and HES1 (green) in primary tumor from melanoma patients 1. Magnification, X40. (J) CCK8 proliferation assay of HES1^low^ senescent CTCs, HES1^high^ senescent CTCs, HES1^low^ senescent CTCs with LCS-1(SOD1 inhibitor, 0.1uM) treatment, and HES1^high^ senescent CTCs with rSOD1 (SOD1 recombinant protein, 50ng/ml) treatment. *n* = 3 independent experiments. (K) Quantification analysis of Mitosox Red in HES1^low^ senescent CTCs, HES1^high^ senescent CTCs, HES1^low^ senescent CTCs with LCS-1(SOD1 inhibitor) treatment, and HES1^high^ senescent CTCs with SOD1 recombinant protein treatment. *n* = 3 independent experiments. (L) Representative images of mitochondrial morphology analysis by TEM in HES1^low^ senescent CTCs, HES1^high^ senescent CTCs, HES1^low^ senescent CTCs with LCS-1(SOD1 inhibitor) treatment, and HES1^high^ senescent CTCs with SOD1 recombinant protein treatment. (M) Quantification of length ratio of cristae to outer mitochondrial membrane, and cristae length of HES1^low^ senescent CTCs, HES1^high^ senescent CTCs, HES1^low^ senescent CTCs with LCS-1 (SOD1 inhibitor) treatment, and HES1^high^ senescent CTCs with SOD1 recombinant protein treatment. (N) Schematic overview of the mechanism underlying HES1-SOD1-ROS axis regulate relapse of senescent CTCs. **B, D, G**, **J, K,** and **M,** Data are represented as mean ± SEM. **B, D, G**, **J, K,** and **M,** Statistical significance was calculated by a two-sided Student *t* test; ***, *P* < 0.001; ****, *P* < 0.0001. RFI, relative fluorescence intensity; ROS, reactive oxygen species.LCS-1, SOD1 inhibitor. rSOD1, SOD1 recombinant protein.

To explore molecular mechanisms mediating mitochondrial homeostasis and ROS balance, we integrated epigenomic CUT&Tag sequencing and single-cell transcriptomic analysis to identify direct downstream targets of HES1 in Mel-167 senescent CTCs. Among the 10 candidate targets identified, the most highly upregulated hit was Superoxide Dismutase 1 (*SOD1*) (Figure 5E). Direct visualization of the CUT&Tag sequencing reads revealed a sharp peak of HES1 binding at the promoter region of *SOD1* (Figure 5F). This raised the possibility that HES1 can directly regulate *SOD1* transcription. Dual-luciferase reporter assays confirmed that HES1 effectively suppressed *SOD1* promoter activity *in vitro* (Figure 5G). Knockout (KO) of HES1 in CTCs led to significant upregulation of the mRNA and protein levels of SOD1 (Figure S5B-C). Conversely, overexpression of HES1 inhibited the expression of SOD1, resulting subsequent elevation of ROS levels *in vitro* (Figure S5D-F). Consistently, a significant increase of SOD1 expression was observed in Sen_HES1^low^ CTCs compared to Sen_HES1^high^ CTCs. (Figure S5G-H).

We further assessed the expression differences of HES1 and SOD1 *in vivo*. The results showed that SOD1 was significantly higher in Sen_HES1^low^ CTC-derived tumors than the Sen_HES1^high^ mice tumors (Figure S5I). Consistently, an antagonistic expression pattern between HES1 and SOD1 protein was also observed in mice tumors derived from these two senescent CTC subpopulations as measured by Multiplex immunohistochemistry (mIHC) staining (Figure 5H; Figure S5J). A similar inverse correlation of HES1 and SOD1 expression was also observed in melanoma clinical samples (Figure 5I). Thus, HES1 can directly repress SOD1 transcription, which may contribute to the antagonistic expression patterns of these two genes.

To test the causal relationship between the HES1-SOD1 axis and mitochondrial functionality, we treated these senescent CTCs with SOD1 inhibitor (LCS-1) or recombinant SOD1 (rSOD1) to monitor the regrowth potential. Interestingly, SOD1 inhibition significantly impaired the regrowth capability of Sen_HES1^low^ CTCs, while the supplementation of recombinant SOD1 protein in Sen_HES1^high^ CTCs substantially enhanced the regrowth of these cells (Figure 5J, Figure S5K). Consistently, SOD1 inhibitor treatment in Sen_HES1^low^ CTCs resulted in elevated ROS levels, while rSOD1 addition significantly reduced ROS accumulation in Sen_HES1^high^ CTCs (Figure 5K; Figure S5L). TEM analysis further showed that SOD1 inhibitor treatment increased aberrant mitochondria structures. Conversely, rSOD1 addition in Sen_HES1^high^ CTCs significantly rescued the mitochondrial structural defects (Figure 5L-M; Figure S5M).

Collectively, these results suggested that HES1 can directly repress SOD1 transcription, subsequently leading to differential expression of HES1 and SOD1 in senescent CTC subpopulations. The antagonism between HES1 and SOD1 impacted mitochondrial functionality and ROS detoxification capability, which likely contributed to the relapse potential of senescent CTCs *in vitro* and *in vivo* (Figure 5N).

### Senescent CTC subpopulations exhibit multi-drug resistance *in vitro*

We next assessed the clinical relevance of these senescent CTC subpopulations. Mel-167 is a polyclonal BRAF^V600E^ mutant melanoma CTC cell line that exhibits differential sensitivities to various cancer-targeting drugs^30^. We treated senescent and non-senescent CTC with classical anti-tumor chemotherapy (MTX and TMZ), BRAF^V600E^-targeted therapy (Vemurafenib, named as BRAFi) and a ferroptosis inducer (Erastin). Remarkably, while non-senescent CTCs showed different degrees of sensitivities to these drugs, both senescent CTC subpopulations demonstrated strong multi-drug resistance phenotypes, suggesting these dangerous tumor subclones might represent a potential clinical challenge (Figure 6A).

**Figure 6.**
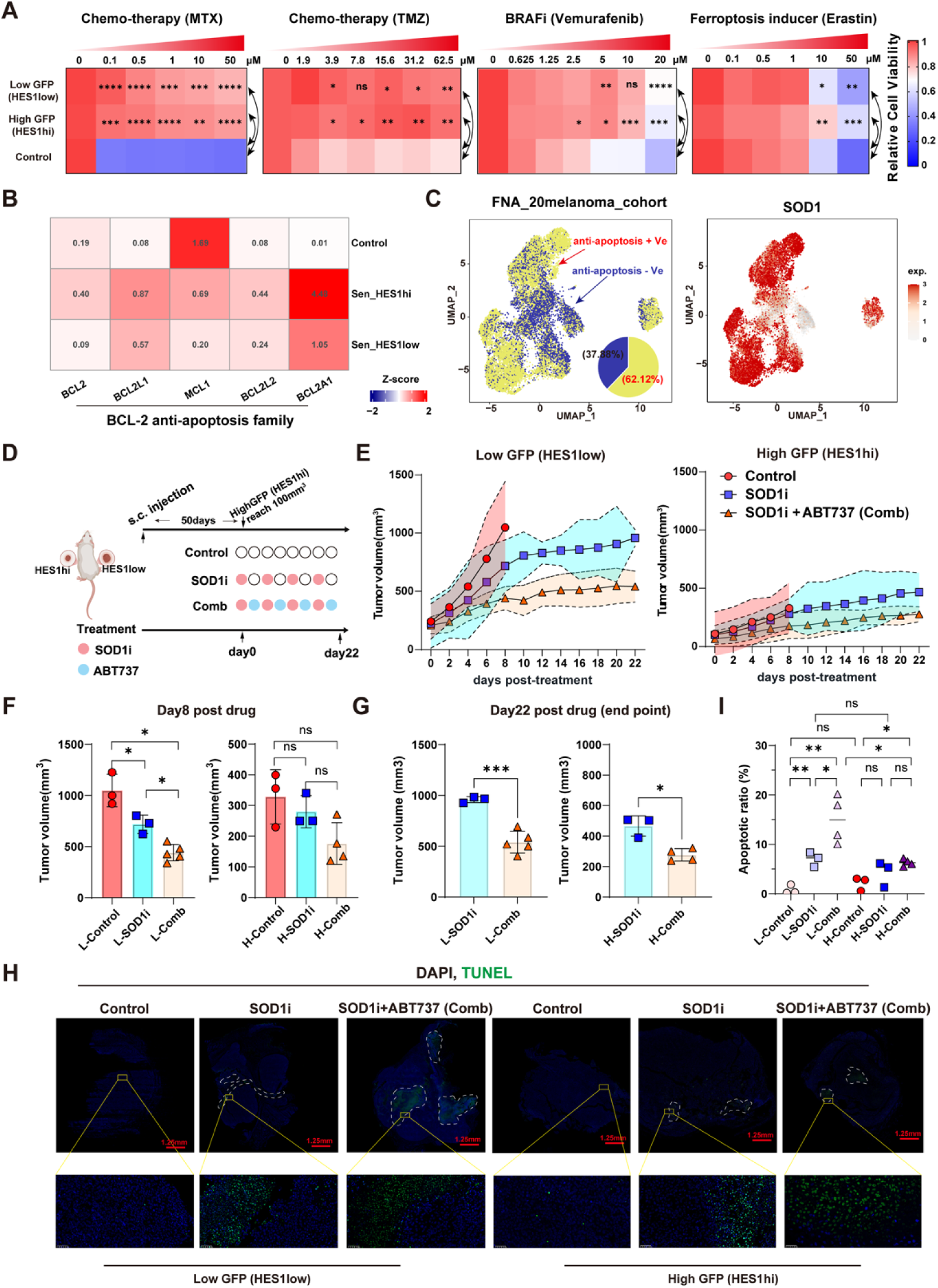
Combination therapy of SOD1 inhibitor and senolytic drug ABT737 suppresses tumor recurrence *in vivo*. (A) Heatmap represents drug sensitivity of methotrexate (MTX), Temozolomide (TMZ), BRAFi (Vemurafenib) and Ferroptosis inducer (Erastin) in senescent and non-senescent CTCs. (B) Heatmap shows the mRNA expression level of BCL-2 family members (*BCL2*, *BCL2L1*, *BCL2A1*, *BCL2L2*, *MCL1*) among Sen_HES1^low^, Sen_HES1^high^ senescent CTC and wild-type CTCs in scRNA-seq. (C) UMAP visualization of anti-apoptosis signatures positive (anti-apoptosis + Ve) melanoma cells in **Figure S6B** (left). Each dot indicates a single cell. Color-coded for BCL-2 family signatures positive or negative. Pie plot shows percentage of anti-apoptosis + Ve melanoma cells among all melanoma cells. UMAP visualization of SOD1 expression in **Figure S6B** (right). (D) Schematic diagram of depicted *in vivo* experiment of subcutaneous injection of HES1^low^ senescent CTCs, HES1^high^ senescent CTCs into NCG mice. Relapse without treatment (Control) group: *n* = 3 mice; Relapse with SOD1 inhibitor (SOD1i) group: *n* = 3 mice; Relapse with SOD1 inhibitor+ ABT737 (Comb) group: *n* = 5 mice. The time of drug treatment was defined when the average tumor volume in HES1^high^ group reached to 100mm^3^ (defined as day 0). SOD1i (LCS-1, 20mg/kg, intraperitoneal injection) on day 0 and thereafter, every two days for three weeks; ABT (ABT737, 30mg/kg, oral gavage) on day 0 and thereafter, every two days for three weeks. (E) Tumor growth curves of Control, SOD1i and Comb groups under drug treatment. (F) Quantification analysis of HES1^low^ and HES1^high^ relapse tumors volume in Control, SOD1i and Comb groups at day8 post drug treatment. L-Control, L-SOD1i, and L-Comb refer to HES1^low^ relapse tumors in Control, SOD1i and Comb groups, respectively. H-Control, H-SOD1i, and H-Comb refer to HES1^high^ relapse tumors in Control, SOD1i and Comb groups, respectively. (G) Quantification analysis of HES1^low^ and HES1^high^ relapse tumors volume in Control, SOD1i and Comb groups at experiment endpoint. (H) Representative images of apoptotic cells via TUNEL staining of HES1^low^ and HES1^high^ relapse tumors in Control, SOD1i and Comb groups. Apoptotic area was circuit with white dash line. Magnification, x4 and x40. (I) Quantification of apoptotic cells via TUNEL staining of relapse tumors from Figure 6H. **F, G,** and **I,** Data are represented as mean ± SEM. **F,** and **I,** Statistical significance was calculated by two-way ANOVA with the Bonferroni *post hoc* test. **A**, and **G,** Statistical significance was calculated by a two-sided Student *t* test. NS, not significant; *, *P* < 0.05; **, *P* < 0.01; ***, *P* < 0.001. TUNEL, TdT-mediated dUTP Nick-End Labeling.

### Combination therapy of SOD1 inhibitor and senolytic drug ABT737 suppresses tumor recurrence *in vivo*

We observed the upregulation of BCL-2 anti-apoptosis family genes and the hyperactivation of apoptosis suppression pathway in Sen_HES1^high^ and Sen_HES1^low^ subpopulations, indicating that senescent CTCs are likely sensitive to apoptosis-inducing senolytic drug ABT737 (Figure 6B; Figure S6A). Interestingly, in an independent analysis of single-cell transcriptomic dataset of melanoma patients whose disease recurred following first-line therapy^31^, we found that anti-apoptotic BCL2 signature-positive tumor cells correlated well with SOD1 positivity, and these cells accounted for 62.12% of the total tumor cell population (Figure 6C; Figure S6B-C).

These data indicated that SOD1 expression was likely linked to anti-apoptosis function.

Given the capability of SOD1 in ROS detoxification which may contribute to tumor regrowth potential, we further tested the combination therapeutic strategy in suppressing tumor recurrence from senescent Mel-167 CTCs. Mice xenograft tumors were generated using pair-wise subcutaneous injection of Sen_HES1^high^ and Sen_HES1^low^ CTCs in each flank near the hind leg of the mice. After 50 days of CTC implantation, the drug treatment was started when the average tumor volume in Sen_HES1^high^ group reached about 100mm^3^ (defined as day 0) in the following groups: (1) control treatment (Control), (2) SOD1i (LCS-1) single drug treatment (starting on day 0, and the drug was given once every two days), and (3) combination treatment with LCS-1+ ABT737 (Comb, starting on day 0, and the two drugs were given at alternative days once every two days). Mice tumor burden was carefully monitored over 22 days of drug treatment. Similar to the results shown in Figure 3C-E, the total tumor burden derived from Sen_HES1^low^ CTCs was significantly higher compared to the Sen_HES1^high^ group at day 0 of drug treatment (Figure 6D; Figure S7A). Control tumors derived from senescent CTCs grew very large and reached the endpoint at day 8 of drug treatment. In contrast, SOD1i treatment group significantly attenuated tumor regrowth in Sen_HES1^low^ group, and the combination therapy achieved an even stronger tumor inhibition when compared to SOD1i treatment group (Figure 6E-G). The anti-tumor activities of SOD1i and ABT737 correlated well with the ability to induce apoptosis, with the combination therapy showing maximal apoptosis induction *in vivo* in the Sen_HES1^low^ group (Figure 6H-I). Similar tumor suppressive effects were observed in Sen_HES1^high^ CTC-derived tumor group treated with SOD1i or combination therapy, although the ability to induce apoptosis was weaker in the Sen_HES1^high^ group when compared to the Sen_HES1^low^ group (Figure 6D-I). Interestingly, when compared to the control group, the anti-proliferation effect (marked by MKI67 IHC staining) was only observed in the combination group, but not the SOD1i treatment group, suggesting SOD1 inhibition alone did not affect tumor cell proliferation *in vivo* (Figure S7B-C).

In summary, SOD1 inhibitor combined with the senolytic drug ABT737 treatment achieved a synergistic suppression of tumor relapse in preclinical mouse models implanted with senescent CTCs.

### Senescent melanoma CTCs are associated with therapeutic resistance

To further investigate the clinical relevance of senescent CTCs in advanced melanoma, a total of 28 on-treatment melanoma patients were prospectively enrolled (Clinical trial NO. NCT07584291), whose baseline characteristics are summarized in Table S1. The majority of patients (27/28) received immunotherapy (Anti-PD1+ IFNα1b), and one patient received targeted therapy (BRAFi + MEKi). Based on standard RECIST 1.1, patients were classified into progressive disease (PD, n=14) and non-progressing disease (non-PD, n=14).

Melanoma CTCs were isolated from whole blood using SEED-X1.0 microfluidic platform, and analyzed by multiplexed immunofluorescence. Senescent CTCs were identified using staining positivity for nucleic DAPI, HMB45, β-gal, and negativity for CD45. The senescent CTCs were further stained for HES1 and SOD1 expressions to monitor distinct subtypes (Figure 7A). In this cohort, 24 patients (85.7%) contain at least one senescent CTC with a median number of 31 per 7.5 milliliter (ml) of whole blood (range: 0-164). The median total CTC number (senescent and non-senescent CTCs) was 67 per 7.5 milliliter (ml) of whole blood (range 8-220) (Table 1; Table S1). Remarkably, PD patients showed significant elevations of β-gal^+^ senescent CTCs than the non-PD group, with at least half of PD patients with > 50% of all CTCs scored positive for β-gal staining, suggesting the dominant presence of senescent CTCs in these patients (Figure 7B, *P* < 0.05). The differences were even more evident for the total abundance and fraction of β-gal^+^ HES1^+^ senescent CTC subtype in PD versus non-PD patients (Figure 7B*, P* < 0.01), while the β-gal^+^ SOD1^+^ senescent CTC subtype did not reach statistical significance between the two groups (Figure 7B). Interestingly, the total number of CTCs (HMB45^+^ and CD45^-^) were also significantly correlated with disease progression, although the non-senescent CTC subtypes, including β-gal- HES1^+^ CTCs and β-gal^-^ SOD1^+^ CTCs, did not show such correlation (Figure 7C; Figure S8A).

**Figure 7.**
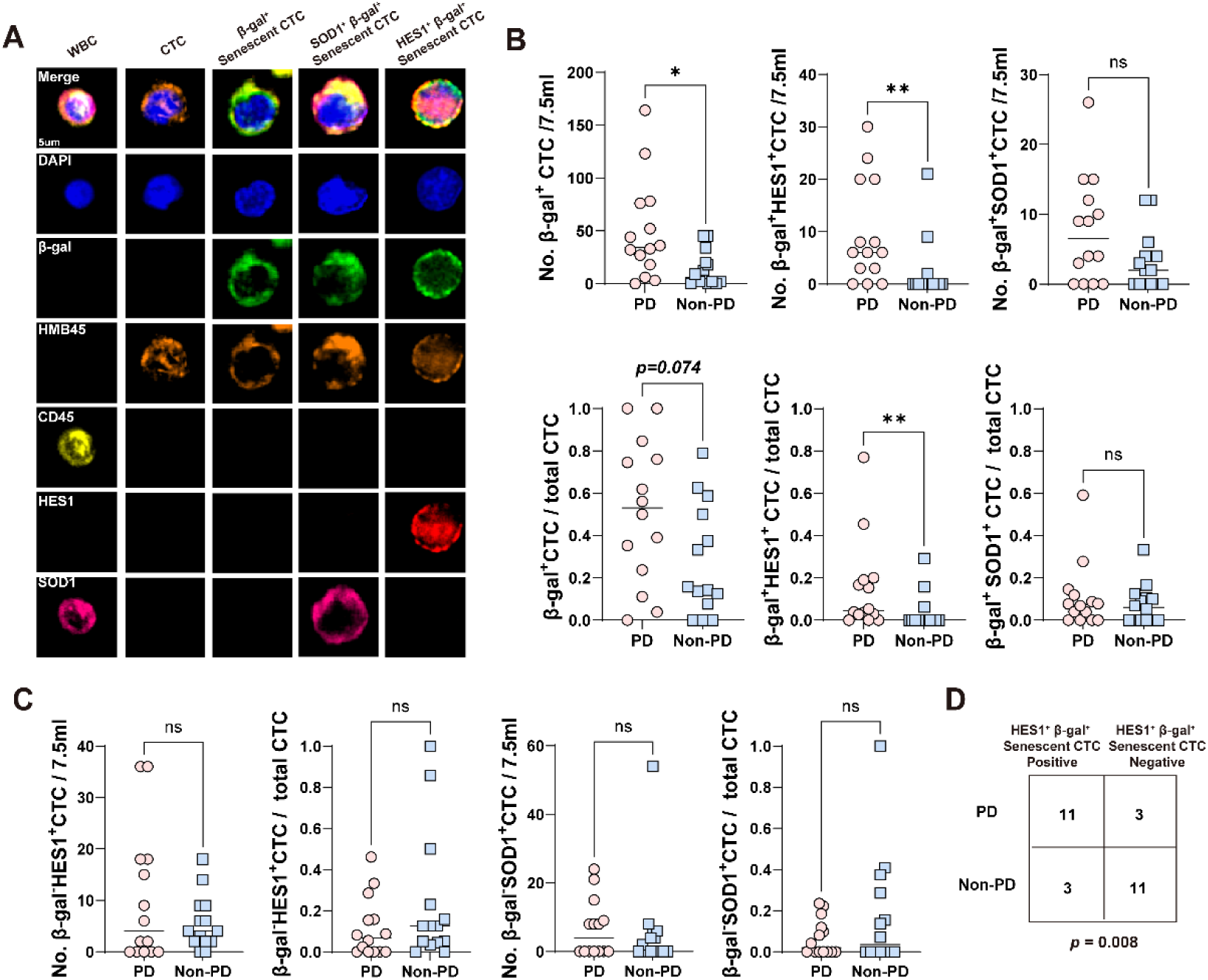
Senescent melanoma CTCs are associated with therapeutic resistance. (A) Representative images of mIF staining of CTCs shows DAPI (blue), β-gal (green), HMB45 (orange), CD45 (yellow), HES1 (red) and SOD1 (pink) in CTCs isolated from first-line on-treatment melanoma patients. HMB45^+^CD45^-^ cells were recognized as CTCs. β-gal^+^HMB45^+^CD45^-^ cells were recognized as senescent CTCs. Scale bar, 5 μm. (B) Box plot (top) showing the number of β-gal^+^ senescent CTCs, β-gal^+^HES1^+^ senescent CTCs and β-gal^+^SOD1^+^ senescent CTCs. Box plot (bottom) showing the proportion of β-gal^+^ senescent CTCs, β-gal^+^HES1^+^ senescent CTCs and β-gal^+^SOD1^+^ senescent CTCs among all CTCs. (PD, progress disease, *n* =14; non-PD, stable disease or partial response, *n* =14). (C) Box plot (left) showing the number of β-gal^-^HES1^+^ CTCs and β-gal^-^SOD1^+^ CTCs. Box plot (right) showing the proportion of β-gal^-^HES1^+^ CTCs and β-gal^-^SOD1^+^ CTCs among all CTCs. (D) Fisher test was performed to assess the association between β-gal^+^HES1^+^ senescent CTCs and clinical outcome. **B,** and **C,** Data are represented as mean ± SEM. **B,** and **C,** Statistical significance was calculated by Mann-Whitney U test; **D,** statistical significance was calculated by Fisher test. NS, not significant; *, *P* < 0.05; **, *P* < 0.01.

**Table 1.**
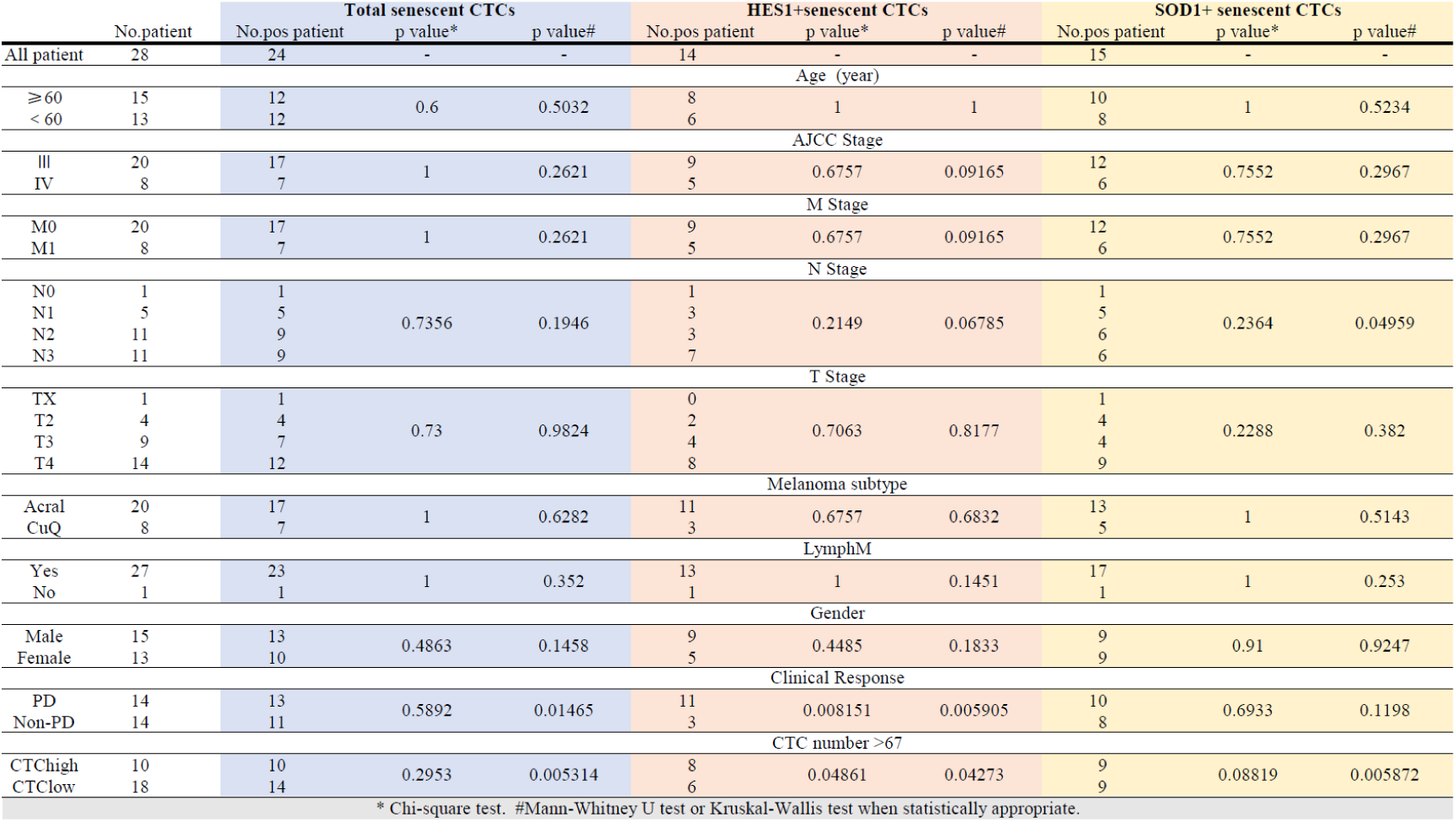
Demographic, patient characteristics, and abundance of total senescent CTC, HES1^+^senescent CTC, and SOD1^+^ senescent CTC. (. No., number of. Cutoff: 1 senescent CTCs per patient)

Collectively, our data revealed the widespread presence of β-gal^+^ senescent CTCs in blood samples of on-treatment melanoma patients, and both the total number and fraction β-gal^+^ HES1^+^ senescent CTCs were strongly associated with disease progression (Figure 7D, Figure S8B). These preliminary findings warrant further clinical validation in larger cohort of melanoma patients.

## DISCUSSION

Conventionally, senescence is thought as a tumor-suppressive mechanism that enforces irreversible proliferative arrest^3–5^. In the present study, by integrating cross-disciplinary technologies including single-cell multi-omics, microfluidic CTC isolation platform, experimental CTC model systems and clinical analyses, we systematically probed molecular and functional heterogeneity of senescent CTCs, and identified two senescent CTC subpopulations marked by HES1 expression levels. Both CTC subpopulations exhibited classical features of senescence, yet they are operated under distinct molecular and metabolic programs, and are functionally linked to differential metastatic regrowth potential. Interestingly, Hu et al. first reported that senescent melanoma CTCs are broadly distributed in patients, which are correlate with treatment failure, suggesting that these cells are not merely bystanders^2^. Similarly, Duy et al. showed that a senescence-like phenotype allows AML cells to survive chemotherapy and subsequently repopulate the leukemia, suggesting that the disruption of senescence-like arrest is a conserved survival strategy across tumor types^13^. In line with this, Zhang et al. found that radiotherapy-induced tumor senescence activates epithelial–mesenchymal transition module in cervical carcinoma, which in turn facilitates disease recurrence^32^. By identifying the specific molecular signatures that mark relapse-competent senescent CTCs, our study suggested that these cells could be clinically dangerous tumor subclones driving metastatic relapse. These findings also highlighted the need to therapeutically target this previously overlooked minimal residual disease in blood before it develops into overt metastasis.

Our work suggested that senescent CTCs are functionally heterogeneous. Sen_HES1^low^ CTCs showed substantially enhanced tumor regrowth and an elevated pro-inflammatory and thrombotic phenotype than the Sen_HES1^high^ CTCs (Figure 3). This difference was accompanied by a striking divergence in mitochondrial functionality: Sen_HES1^low^ CTCs maintained higher membrane potential, oxidative respiration, and mitophagic activity, whereas Sen_HES1^high^ CTCs accumulated structurally and functionally abnormal mitochondria and elevated intracellular ROS levels, indicative of severely impaired oxidative metabolism (Figure 4). Multiple studies across different cancers have demonstrated a causal link of mitochondrial dysfunction to tumor senescence^3,33,34^. Our findings implied that the capability of senescent tumor cells to preserve mitochondrial functionality might be directly linked the tumor reactivation potential from senescent states. Thus, functional perturbation of mitochondrial-associated programs in experimental models may enable us to identify novel factors involved in metastatic relapse.

At the mechanistic level, our data identified direct transcriptional repression of SOD1 by HES1, as a molecular determinant linking distinct CTC states to ROS detoxification and tumor regrowth competence. The antagonism between HES1 and SOD1 induced functionally distinct CTC senescence states. This finding raised the likelihood that heterogenous senescent subclones may co-exist and cooperate with each other to drive disease progression rather than in an isolated clonal state. The molecular crosstalk among distinct tumor subpopulations has been documented in several studies. For example, Ma et al reported that CCF^+^ (cytoplasmic chromatin fragments) senescent tumor cells secrete CCF following chemotherapy to promote stemness of CCF^-^ group to facilitate tumor recurrence^8^. Goyette et al reported that in tumors heterogeneous for HER2 expression, HER2^low^ subclones support the tumorigenic fitness and targeted therapy resistance of HER2^hi^ cells^35^. Whether Sen_HES1^low^ and Sen_HES1^high^ CTCs can similarly communicate with each other within the pre-metastatic niches warrants further investigation.

Critically, our work showed that despite the differences in tumor regrowth potential of these senescent CTC subpopulations (Figure 3), both groups share the traits of multi-drug resistance to cytotoxic and targeted agents (Figure 6). This highlighted that the therapeutic challenge posed by senescent CTCs extends across different subgroups. An important observation of this study is that the upregulation of BCL-2 anti-apoptosis family members in senescent CTCs contributed to tumor relapsed *in vivo* (Figure 6A-B; Figure S6A). Consistent with our observation, recurrent tumors from melanoma patients after first-line treatment also exhibited elevated anti-apoptosis program at single-cell resolution, suggesting a crucial role of the BCL-2 anti-apoptosis family members in tumor recurrence (Figure S6B-D)^31^. These observations provided a rationale for combining redox disruption with senolytic targeting to eliminate senescent CTCs. Indeed, our proof-of-concept preclinical testing combining SOD1i (LCS-1) and the anti-apoptosis senolytic drug ABT737 suppressed senescent CTC-derived tumor growth more effectively than SOD1i treatment alone (Figure 6). Consistently, emerging studies support BCL-2 family dependence as a recurrent survival mechanism in stem-like and therapy-resistant tumor states^36–39^, and BH3-mimetic approaches can restore drug sensitivity in preclinical models of neuroblastoma^40^, lung cancer^41^, ovarian carcinoma^42^, melanoma^37^, and rhabdomyosarcoma^43^. Clinically, the BCL-2 inhibitor venetoclax has demonstrated activity in relapsed hematological malignancies^44–46^. Nevertheless, it remains to be established whether such combination therapeutic strategy may achieve clinical benefit in recurrent melanoma and other cancer types.

The clinical data underscored the potential value of resolving distinct CTC states, rather than relying on total CTC enumeration alone. Molecularly defined CTC subsets have increasingly shown powerful prognostic biomarker values ^1,47,48^, including DLL3⁺ CTCs in small-cell lung cancer^49^ and GPNMB⁺ CTCs in brain metastatic disease^50^. In our prospective on-treatment melanoma cohort, the abundance of β-gal⁺ senescent CTCs, and the number and fraction of β-gal⁺HES1⁺ CTCs were significantly higher in patients with progressive disease, supporting their evaluation as a candidate response-monitoring biomarker (Figure 7). We did not found a significant association of β-gal⁺SOD1⁺ CTCs with disease progression. One possibility is that β-gal⁺HES1⁺ CTCs mark persistence or accumulation of a treatment-resistant senescent reservoir rather than the per-cell probability of subsequent tumor reactivation. Longitudinal sampling with matched clinical outcomes would be required to determine whether dynamic changes in these distinct CTC states predict response and metastatic relapse.

In summary, our study has uncovered substantial CTC senescence heterogeneity driven by distinct molecular and metabolic programs. Furthermore, we identified the antagonistic regulation between HES1 and SOD1 that controlled mitochondrial fitness and ROS homeostasis, which contributed to CTC regrowth potential. Preclinical evidence suggested that combination treatment using SOD1i and ABT737 could effectively suppress senescent CTC-induced tumor relapse *in vivo*. In a prospective analysis of melanoma patient samples, the prognostic impact of β-gal⁺HES1^+^ senescent CTCs on therapeutic resistance highlighted their potential for noninvasive disease monitoring. Nevertheless, our clinical analysis contained limited number of patients and multi-center studies with larger patient cohorts are needed to validate these findings.

## METHODS

### Patient cohort and blood sample collection

This prospective, observational cohort study aims to explore the multi-omics profiles of liquid biopsies and develop clinical biomarkers in melanoma (ClinicalTrials.gov Identifier: <u>NCT07584291</u>). Fresh peripheral blood samples (approximately 7-10 mL) were collected from patients diagnosed with melanoma at Xijing Hospital, Fourth Military Medical University between Nov 11, 2025 and July 15, 2026. Clinical response criteria were assessed as per RECIST version 1.1. This study was conducted in accordance with the principles of the Declaration of Helsinki and was approved by the Institutional Ethics Review Committee of Xijing Hospital (Protocol No. KY20252289-C-1). Prior to enrollment and blood sample collection, all participants provided written informed consent after receiving a detailed explanation of the study objectives and procedures. Clinical staging of melanoma was assessed according to the 9th edition of the tumor–node–metastasis (TNM) classification system. Peripheral blood samples were collected using sterile anticoagulant-containing tubes and were processed immediately for downstream experiments to ensure optimal preservation of cell viability and biological activity. Detailed clinicopathological characteristics of all enrolled patients, including age, sex, and TNM stage, are summarized in Supplementary **Table S1**.

### CTC isolation

The isolation of circulating tumor cells (CTCs) from the peripheral blood of skin melanoma patients was performed utilizing the SEED-X1.0, a novel inertial microfluidic platform designed for ultra-fast, label-free cell enrichment. The separation mechanism utilizes a four-stage curved channel design that leverages the synergistic effects of inertial lift forces and Dean drag forces to isolate target cells based on size. As the sample traverses the spiral channels, larger cells such as melanoma CTCs—typically ranging from 15 to 25 µm—are focused toward the inner wall of the elutriation subchannels, while smaller hematocytes are progressively diverted into waste subchannels. This approach processes whole blood directly without the need for antibody labeling or erythrocyte lysis, which minimizes cell damage and prevents the loss of rare target cells. Under an optimized total flow rate of 10.9 mL/min, the device can process 5-7 mL of whole blood in less than 10 minutes. This high-throughput workflow achieves a white blood cell depletion rate of 99.95%, allowing for the efficient recovery of CTCs for downstream immunofluorescence staining.

### Cell lines and Cell culture

The Mel-167 circulating tumor cell line was established in our previous work from the peripheral blood of a melanoma patient and successfully developed into a stable, expandable in vitro cell line^30^. The Mel-167 was cultured in RPMI-1640 medium (Gibco, USA) supplemented with 20 ng/mL EGF (Gibco, USA), 20 ng/mL bFGF (Gibco, USA), and 10 mL of 1× B27 supplement (Gibco, USA) per 500 mL of medium, under hypoxic conditions (4 % O₂) with 5 % CO₂ at 37 °C. The 293T cell line (ATCC) was maintained in high-glucose DMEM (Gibco, USA) supplemented with 10 % fetal bovine serum (FBS; ExCell Bio, China) under standard culture conditions (5 % CO₂, 37 °C). All cell lines were routinely monitored for morphology and viability, screened for mycoplasma contamination, and used within a limited number of passages to ensure reproducibility and experimental consistency.

### Ethics approval and consent to participate

All samples for CTC mIF were collected at Fourth Military Medical University Xijing Hospital (approval number: KY20252289-C-1). All mouse experiments were approved by the Institutional Animal Care and Use Committee of SUSTECH (approval number: SUSTech-JY202402014).

### Antibodies, plasmids, primers, and short hairpin RNA/siRNA

Information on antibodies, chemicals, primer sequences, short hairpin RNAs (shRNA), and siRNAs can be found in Supplementary Tables S2–S4.

### Lentivirus preparation

In our previous work, doxycycline (DOX)-inducible CTTN knockdown Mel-167 cells were successfully established in our laboratory^2^. Building on this cell line, the cells were subsequently transduced with lentivirus encoding the HES1 promoter-GFP construct for downstream experiments. Briefly, the HES1 promoter sequence was synthesized and inserted into the Fused pSin-P2A-GFP plasmid vector, which had been linearized using BamHI-HF (NEB, R3136T) and AgeI-HF (NEB, R3552S) restriction endonucleases, ensuring that the HES1 promoter was positioned immediately upstream of the GFP coding sequence. For lentiviral production, 7 µg of the resulting vector (Fused pSin-P2A-HES1 promoter-GFP) was co-transfected with 7 µg of psPAX2 (RRID: Addgene_12260) and 2.4 µg of pMD2.G (RRID: Addgene_12259) into HEK293T cells using polyethylenimine as the transfection reagent. Sixty hours post-transfection, the lentiviral supernatants were collected and filtered through a 0.45 µm syringe filter (Thermo Fisher 7232545) to remove cell debris. For target cell infection, 0.3–0.75 ml of the lentiviral supernatant was added to Mel-167 cells in the presence of 1.5 µg/ml polybrene and incubated for 16 h in low-attachment 6-well plates to enhance transduction efficiency. Following infection, GFP-positive cells were enriched and isolated by flow cytometry for downstream experiments.

### Flow Cytometric Isolation of HES1^high^ and HES1^low^ Mel-167 Subpopulations

Based on bioinformatic analysis, heterogeneity in HES1 expression was observed in Mel-167 cells following CTTN knockdown. To isolate subpopulations with high and low HES1 expression, DOX-inducible CTTN knockdown Mel-167 cells were further transduced with a lentiviral construct encoding the HES1 promoter-driven GFP reporter. GFP fluorescence intensity was used to distinguish and sort cells with high and low HES1 expression. Briefly, 48 hours after DOX induction, cells were harvested, centrifuged, and resuspended in phosphate-buffered saline (PBS). GFP-high and GFP-low subpopulations were sorted using a FACSAria II flow cytometer (BD Biosciences, USA), collected separately, centrifuged to remove supernatant, and resuspended in CTC culture medium. Cells were allowed to recover in a culture incubator before downstream experiments. Sorting gates were defined based on negative controls (non-transduced cells) to accurately distinguish high and low GFP-expressing subpopulations.

### Cell Proliferation Assay

GFP-high and GFP-low Mel-167 subpopulations, isolated by flow cytometry, were used to assess cell proliferation in both 2D and 3D culture systems. For 2D proliferation, cells were seeded in 96-well plates at a density of 5 × 10³ cells per well. Cell viability was measured at 0, 5, and 10 days using the Cell Counting Kit-8 (GLPBIO, GK10001) according to the manufacturer’s instructions. Absorbance at 450 nm was recorded using a microplate reader. For 3D proliferation, cells were embedded in ultra-low attachment 96-well plates to form spheroids, with the same seeding density, and cultured under standard conditions. Cell viability of spheroids was similarly assessed at 0, 5, and 10 days using CellTiter-Glo® 3D Cell Viability Assay Kit (Promega, G9682) following the product manual, and absorbance values were recorded. Relative cell viability was calculated by normalizing the absorbance at each time point to the initial measurement. All experiments were performed in triplicate.

### Immunofluorescence Analysis

Sorted Mel-167 cells (GFP-high and GFP-low subpopulations), isolated by flow cytometry following DOX-induced CTTN knockdown, were subjected to immunofluorescence staining to assess HES1 expression. Cells were spun onto adherent glass slides using a cytospin, then fixed with 4 % paraformaldehyde (PFA) for 15 min at room temperature. After washing twice with PBS, cells were permeabilized with 0.5 % Triton X-100 for 10 min. Cells were then blocked with 5 % bovine serum albumin (BSA) for 1 h at room temperature and incubated with a primary antibody Lysosome marker LAMP1 (CST, 42406T, dilution 1:200), and mitochondrial marker TOM20 (CST, 156655S, dilution 1:400) at 4 °C overnight.

Following washes with TBST, cells were incubated with a fluorescently labeled secondary antibody (1:1000) for 1 h at room temperature in the dark. Nuclei were counterstained with 1 μM Hoechst 33342 (Invitrogen, H3570) for 5 min. Fluorescence images were captured using a confocal microscope.

### Immunoblotting

Cells were lysed in RIPA buffer supplemented with proteinase inhibitor cocktail (Roche, 04693132001) and phosphatase inhibitor cocktails (Roche, 04906837001) on ice for 30 min. The lysates were centrifuged at 12,000 × g for 15 min at 4°C, and the supernatants were collected for protein concentration determination using a BCA Protein Assay Kit (Pierce, 23225). Equal amounts of protein were mixed with loading buffer, boiled at 95°C for 5 min, separated by SDS-PAGE, and transferred onto PVDF membranes. The membranes were blocked with 5% non-fat milk for 1 h at room temperature and then incubated with the indicated primary antibodies overnight at 4°C. After washing with TBST, the membranes were incubated with HRP-conjugated secondary antibodies for 1 h at room temperature. Protein bands were visualized using an enhanced chemiluminescence (ECL) detection Kit (Pierce, 32106). GAPDH or Tubulin was used as a loading control.

### RT-qPCR

Total RNA was extracted using the NucleoZOL reagent (Macherey-Nagel, 74040.200) according to the product manual. RT was performed using the EvoM-MLV reverse transcription Kit (Accurate Biology, AG11728) following the product manual. qPCR was prepared using the SYBR Green premix pro Taq HS Kit (Accurate Biology, AG11718), and then qPCR was performed using a QuantStudio 12K Flex Real-Time PCR System (Applied Biosystems, RRID:SCR_021098).

### In Vivo Tumor Growth and Metastasis Assays

GFP-high and GFP-low Mel-167 subpopulations, isolated by flow cytometry, were used for in vivo experiments. All animal studies were approved by the Institutional

Animal Care and Use Committee of the Southern University of Science and Technology (IACUC NO. SUSTech-JY202402014). NCG mice (6–8 weeks old) were purchased from Gempharmatech Co., Ltd, China. For subcutaneous tumor growth, 8 × 10⁴ cells of each subpopulation were injected subcutaneously into the flanks of NCG mice. Tumor volume was measured every 3–7 days using calipers and calculated using the formula V = (W² × L)/2, where W is tumor width and L is tumor length. At the experimental endpoint, tumors were harvested, weighed, and used for subsequent analyses.

For metastasis assessment, 8 × 10⁴ cells of each subpopulation were injected via the tail vein. Tumor-specific luciferase signals were monitored using an IVIS In Vivo Imaging System (PerkinElmer) to track metastatic burden. All mice were housed under standard conditions with ad libitum access to food and water. Each experiment included at least five mice per group to ensure reproducibility.

### ROS Detection in Tumor Tissue

Fresh subcutaneous tumor tissues derived from GFP-high and GFP-low groups were collected and processed for reactive oxygen species (ROS) detection. This analysis was conducted by Wuhan Pinofi Biotechnology Co., Ltd. Tissue samples were immediately embedded in optimal cutting temperature (OCT) compound and frozen. Cryosections (5–10 µm) were prepared using a cryostat and mounted onto glass slides. Sections were stained using a ROS detection kit (Beyotime, S0063) according to the manufacturer’s instructions. Briefly, sections were incubated with the ROS-sensitive fluorescent probe at the recommended concentration for the specified time at room temperature, protected from light. After washing with phosphate-buffered saline (PBS), sections were counterstained with DAPI to visualize nuclei. Fluorescence images were captured using a confocal microscope, and ROS signal intensity was quantified.

### Transmission Electron Microscopy (TEM) Analysis of Mitochondria

GFP-high and GFP-low, GFP-high treated with recombinant SOD1 and GFP-low treated with LCS-1 (SOD1 inhibitor) Mel-167 subpopulations, isolated by flow cytometry, were used for ultrastructural analysis of mitochondria. This analysis was conducted by Wuhan Servicebio Technology Co., Ltd. Briefly, cells were collected and fixed in TEM fixative (Servicebio, G1102) at 4 °C overnight, followed by post-fixation with 1 % osmium acid in the dark for 2 h. Samples were dehydrated through a graded ethanol series and embedded in EMbed 812 resin (SPI, 90529-77-4). Ultrathin sections (∼60 nm) were prepared using an ultramicrotome (Leica, UC7) and stained with 2 % uranyl acetate in absolute ethanol for 8 min, followed by 2.6 % lead citrate for 8 min. Mitochondrial morphology was examined using a transmission electron microscope (Hitachi, HT7700). For in vivo analysis, tumor tissues derived from GFP-high and GFP-low Mel-167 cells were harvested, cut into ∼1 mm³ pieces, and processed using the same fixation, dehydration, embedding, sectioning, and staining protocol as described for cultured cells. TEM images were acquired to compare mitochondrial ultrastructure between the two subpopulations.

### Drug Sensitivity Assays

GFP-high and GFP-low Mel-167 subpopulations were seeded at a density of 8,000 cells per well in 96-well plates. Cells were then treated with varying concentrations of drugs. Treatments were maintained under standard culture conditions, and cell viability was assessed on day 10 using the Cell Counting Kit-8 (GLPBIO, GK10001) according to the manufacturer’s instructions. Absorbance at 450 nm was measured using a microplate reader, and relative cell viability was calculated by normalizing to untreated controls. All experiments were performed in triplicate to ensure reproducibility.

### Immunohistochemistry

Formalin-fixed, paraffin-embedded (FFPE) tissue blocks from patients or mice were sectioned at a thickness of 5 mm using a microtome (Servicebio, PM-24). For deparaffinization and rehydration, sections were sequentially immersed in three changes of dewaxing solution (G1128) for 10 min each, followed by three changes of absolute ethanol for 5 min each, and finally rinsed with distilled water. Antigen retrieval was performed according to the protocol detailed in the table above. During this process, evaporation of the retrieval buffer was carefully prevented, and sections were not allowed to dry out. After natural cooling, slides were washed with PBS (pH 7.4) on a shaker three times for 5 min each. Endogenous peroxidase activity was quenched by incubating sections in 3% H₂O₂ in methanol for 25 min at room temperature in the dark, followed by three washes in PBS (pH 7.4) for 5 min each on a shaker. Non-specific binding was blocked by applying 3% BSA within the circles drawn by a hydrophobic pen to completely cover the tissue sections for 30 min at room temperature. After blocking, the solution was gently flicked off, and sections were incubated with primary antibodies (anti-CD42b, S0B2175) diluted 1:1000 in PBS overnight at 4°C in a humidified chamber. Following three 5-min washes with PBS (pH 7.4) on a shaker, sections were incubated with HRP-conjugated secondary antibodies applied within the circles for 50 min at room temperature. After three additional washes in PBS (pH 7.4) for 5 min each on a shaker, immunoreactivity was detected using freshly prepared DAB (3,3’-Diaminobenzidine) substrate solution. The color reaction was monitored under a microscope until a positive brownish-yellow signal developed, and was terminated by rinsing with running tap water. Sections were then counterstained with hematoxylin for approximately 3 min, rinsed with tap water, differentiated in acid-alcohol solution for a few seconds, rinsed again with tap water, treated with bluing solution, and washed under running tap water. For dehydration and mounting, two procedures were adopted depending on the sample type. For tissue sections and smears, slides were sequentially dehydrated through 75% ethanol, 85% ethanol, absolute ethanol I, absolute ethanol II, n-butanol, and xylene for 5 min each, briefly air-dried, and mounted with mounting medium. For cells grown on coverslips, coverslips were dehydrated through 75% ethanol, 85% ethanol, absolute ethanol I, and absolute ethanol II for 5 min each, dried with a hairdryer, and mounted cell-side down onto glass slides using neutral balsam. Staining intensity and distribution was analyzed via Saiviewer.

### Double luciferase assay

To evaluate the transcriptional regulation of SOD1 by HES1, HEK293T cells were seeded in 6-well plates at a density of 3 × 10⁵ cells per well and allowed to attach overnight to reach 70–80% confluence at the time of transfection. Cells were co-transfected with the SOD1 promoter luciferase reporter plasmid (firefly luciferase) and either the HES1 overexpression plasmid or the corresponding empty vector, together with a Renilla luciferase plasmid as an internal control, using Polyethylenimine (PEI) as the transfection reagent. Following 48 h of incubation under standard culture conditions (37°C, 5% CO₂), cells were washed with PBS and lysed in passive lysis buffer. Cell lysates were collected and centrifuged to remove debris, and the supernatant was used for luciferase activity measurement. Firefly and Renilla luciferase activities were measured sequentially using the Dual-Luciferase Reporter Assay System on a luminometer, following the manufacturer’s instructions. Firefly luciferase activity was normalized to Renilla luciferase activity, and relative luciferase activity was calculated to quantify the effect of HES1 on SOD1 promoter activity. All experiments were performed in triplicate, and data were expressed as mean ± standard deviation. Appropriate controls, including empty vector and untreated cells, were included in each experiment to ensure reproducibility and accuracy of the results.

### Mitochondrial DNA content analysis

Genomic DNA of cells was isolated using the QIAamp DNA Mini Kit (QIAGEN, 51304). qPCR was prepared using the SYBR Green premix pro Taq HS Kit (Accurate Biology, AG11718), and then qPCR was performed using a QuantStudio 12K Flex Real-Time PCR System. Primers for mitochondrial DNA (mtDNA) tRNAleu (UUR), mtDNA 16S rRNA, and nuclear-encoded B2M were designed. Relative mtDNA content was determined by the ratio of mtDNA tRNAleu (UUR) or mtDNA 16S rRNA to nuclear-encoded B2M.

### Cell-cycle analysis

Cells were collected and washed with PBS once, followed by fixation using precold 75% ethanol at -20°C overnight. After fixation, cells were spun down and washed with PBS and then incubated with the propidium iodide (PI) staining buffer containing 0.1% NP-40, 10 μg/mL RNase (Takara Bio, 2158), and 1:500 PI (Invitrogen, P3566) in PBS for 30 minutes in the dark. Flow cytometry (BD Accuri C6 Flow Cytometer) was used to detect the PI signal using the PE-A channel. The cell cycle was analyzed using FlowJo software.

### Senescence β-galactosidase staining assay

The β-gal staining assay was performed according to the kit instructions (Biosharp, BL133A). MEL-167 cells were treated with or without 300 ng/mL doxycycline for 6 days, fixed with Fixative Solution buffer, washed with PBS, and then incubated overnight in the dark at 37°C with β-gal staining solution. Pictures were captured and analyzed using ImageJ software.

### Mitochondrial membrane potential analysis (JC-1)

JC-1 dye (Invitrogen, T3168) was used to detect the mitochondrial membrane potential. Briefly, cells were collected and washed with Hank’s Balanced Salt Solution (HBSS) buffer (with Ca2^+^, Mg2^+^) and incubated with 2 μmol/L JC-1 in an incubator for 25 minutes. Cells were washed with HBSS buffer and analyzed by flow cytometry using the FL1 and FL2 channels.

### Mitochondrial superoxide analysis

MitoSOX Red (Molecular Probes, M36008) was used to detect mitochondrial superoxide according to the product manual. Briefly, senescent HES1-High and HES1-Low Mel-167 subpopulations in suspension were spun down and washed with HBSS buffer (with Ca2^+^, Mg2^+^), then incubated with 1 μmol/L MitoSOX Red for 30 minutes in a 37°C incubator and washed before flow cytometry analysis.

### Multiplex Immunohistochemistry (mIHC) and HALO analysis

Multiplex immunohistochemistry was performed by a tyramide signal amplification (TSA) six-color kit (abs50014-100T; Absinbio, Shanghai). Tissue slides underwent incubation at 65℃ for 2 hours to enhance adhesion, followed by deparaffinization in xylene and rehydration through an ethanol gradient (100%,95%,70%, 50%). The sections were fixed in 10% neutral buffered formalin for 30 minutes before antigen retrieval using EDTA buffer (pH 9.0, ZSGB-Bio, Beijing, China) under microwave heating. After blocking to reduce nonspecific binding, primary antibodies were applied sequentially alongside horseradish peroxidase (HRP)-conjugated secondary antibodies. Tyramide signal amplification (TSA) was implemented after each primary antibody step to improve detection sensitivity. The sections were then treated with biotinylated rabbit polyclonal anti-rabbit and rabbit anti-mouse secondary antibodies, followed by HRP-conjugated streptavidin as specified by the manufacturer (Absin, abs50014-100T). Chromogenic reactions utilized biotinylated secondary antibodies with streptavidin-linked alkaline phosphatase, while streptavidin-conjugated fluorophores (excitation wavelengths: 520, 570,620, 650, or 780 nm) enabled immunofluorescence detection. Multispectral imaging was conducted using the Vectra Polaris Automated Quantitative Pathology Imaging System (Akoya Biosciences, Delaware, USA).

Positive staining cells were subsequently identified, and advanced spatial analysis was conducted utilizing the HALO™ digital pathology platform (Indica Labs, Corrales, NM, USA). The multiplex immunohistochemistry (IHC) module, which incorporates color deconvolution and nuclear segmentation, facilitated the identification of specific cell subpopulations.

### mIF staining

Peripheral blood samples were collected from patients from XIJING hospital (XIAN, China), CTCs were enriched by the Celutriator TX1 microfluidic device (Shenzhen Genflow Technologies), according to the manufacturer’s operating parameters. The sorted CTCs were fixed with 4% paraformaldehyde (PFA) for 15 min at room temperature (RT). Subsequently, cells were resuspended in PBS and deposited onto positively charged glass slides using a cytocentrifuge (Cytospin™, Thermo Fisher

Scientific) at 1000 rpm for 5 minutes. Cellular senescence was assessed using the CellEvent™ Senescence Green Detection Kit (Thermo Fisher Scientific, C10850). Briefly, the fixed cells on slides were incubated with the CellEvent™ Senescence Green probe solution in a humidified chamber for 1 hour at 37°C, protected from light. Excess reagent was washed off with PBS, and slides were prepared for subsequent staining steps.

mIF was performed by a tyramide signal amplification (TSA) six-color kit (abs50014-100T; Absinbio, Shanghai) and blocked with TBST containing 5% goat serum before incubation with antibodies. To clearly distinguish CTCs from blood cells, we assess the expression of melanoma-associated antigen HMB45 (Dako, M0634, dilution 1:500), leukocyte marker CD45 (Abcam, AB8216, dilution 1:1000), and cortactin (Abcam, AB81208, dilution 1:1500). A cocktail of primary antibodies was prepared in blocking buffer and applied to the slides for 1 hour at RT. Following three washes in PBS, slides were probed with HRP-conjugated anti-rabbit or anti-mouse IgG at RT for 10 min and reacted with fluorophore-conjugated tyramine molecules (PPD 570, PPD 620 or PPD650) for 10 min. At the end of each cycle of staining, the primary and secondary antibodies were washed with eluent solution (absin, abs994) for 10 min at 37°C, respectively. The nuclei were stained with DAPI for 5 min before sealing, and all sections were scanned by a Fluorescent Digital Pathology Slide Scanner (KFBIO, KF-FL-120) at 20× or 40× magnification to generate high-resolution whole slide images (WSIs). The WSIs were subsequently analyzed using PanoScore software (specify version). CTCs were rigorously identified based on the following immunophenotypic criteria: DAPI-positive for an intact nucleus, HMB45-positive, and CD45-negative (DAPI+/HMB45+/CD45-). Leukocytes were identified as DAPI+/CD45+ cells. A CTC was classified as senescent if its cytoplasm exhibited a positive signal from the CellEvent™ Senescence Green probe, co-localizing with the cell’s DAPI and HMB45 signals.

### Hematoxylin and eosin (H&E) staining

Hematoxylin and eosin (H&E) staining was conducted in accordance with established histological protocols to evaluate overall tissue morphology. In summary, formalin-fixed, paraffin-embedded tissue sections underwent deparaffinization in xylene and were subsequently rehydrated through a graded series of alcohol solutions. The sections were then stained with hematoxylin for a duration of two minutes to visualize cell nuclei, followed by differentiation and bluing steps to enhance nuclear contrast. Subsequently, the sections were counterstained with eosin for approximately one to two minutes to delineate cytoplasmic and extracellular matrix components, thereby providing distinct contrast among various tissue structures. Following staining, the sections were dehydrated through a graded alcohol series and cleared in xylene. Finally, the slides were mounted using Entellan® mounting medium (Merck, Darmstadt, Germany) and allowed to dry prior to examination under a light microscope. The stained sections were assessed for architectural features, cellular morphology, and potential pathological alterations.

### Sample preparation and scRNA sequencing

#### Cell preparation

After harvesting, tissues were washed in ice-cold RPMI1640 and dissociated using Multi Tissue Dissociation kit 2 (Miltenyi 130-110-203) from Miltenyi Biotec according to manufacturer’s instructions. DNase treatment was optional according to the viscosity of the homogenate. After erythrocytes removal (Miltenyi 130-094-183), cell number and viability were estimated using Fluorescence Cell Analyzer (Countstar® Rigel S2) with AO/PI reagent, then debris and dead cell depletion (Miltenyi 130-109-398/130-090-101) was determined on live cell results. Finally, fresh cells were washed twice in the RPMI1640 and then resuspended in 1×PBS and 0.04% bovine serum albumin at a concentration of 1×10 6 cells per mL.

#### Single cell Full-length RNA Sequence Transcriptome -seq (scFAST-seq) Library construction and sequencing

scFAST-seq library were prepared using SeekOne® Single Cell Whole Trancriptome Kit according to manufacturer’s instructions (SeekGene Catalog No.K00801). Briefly, an appropriate number of cells were mixed with reverse transcription reagents and added to the sample wells of the SeekOne® DD Chip S3 (Chip S3). Then, Barcoded Hydrogel Beads (BHBs) and partitioning oil were dispensed into corresponding wells separately in Chip S3. Subsequently, Cell-containing reverse transcription reagents and BHBs were encapsulated into emulsion droplets using SeekOne® Digital Droplet System. Immediately following transferring emulsion droplets into PCR tubes, fifteen cycles of annealing (ramping from 8 °C to 42 °C) followed by a 5-min heat inactivation at 85°C were performed to obtain barcoded cDNA. Next, the barcoded cDNA was purified from broken droplet and then twice PCR reactions were performed to remove the majority of ribosomal and mitochondrial cDNA. AMPure beads were used to purify cDNA from the post PCR reaction mixture. Finally, one forth volume of cDNA was fragmented, end repaired, A-tailed and ligated into sequencing adaptor. DNA amplified by index PCR contains any part of polyA or non-PolyA RNA as well as Cell Barcode and Unique Molecular Index. The indexed sequencing libraries were purified using AMPure beads and quantified by quantitative PCR (KAPA Biosystems KK4824). The libraries were then sequenced on Illumina NovaSeq 6000 with PE150 read length or DNBSEQ-T7 platform with PE150 read length.

### Processing the single cell Full-length RNA sequencing data

We used the SeekSoul Tools pipeline to process the cleaned reads and generated the transcript expression matrix. Firstly, the cell barcodes and UMI sequences were extracted based on the defined pattern about the localization of the barcode, linker and UMI within a read. The barcode was corrected with whitelist. The corrected barcode, together with UMI, were put in the header of their corresponding reads. Secondly, the reads were mapped to the reference genomes using STAR 2.5.1b^51^. Then, the reads with barcode and UMI information were assigned to transcriptome using featureCounts of package Subread 1.6.4^52^ (Liao, Smyth et al. 2014). Parameters “ -s ” and “ -t ” of featureCounts vary with different type of chemistries and regions. For the parameter “-s”, “-s 1” was used for product with 3’ chemistry and “-s 2” was used for product with 5’ chemistry. Another parameter “-t exon” was used for read counting only with exon, and “-t transcript” was used for read counting with exon and intron. we also set the parameter “ -fracOverlap ” to 0.5. Other parameters remain default. Finally, similar to the raw_feature_bc_matrix results of Cell Ranger^53^, the raw UMI count matrix according to barcodes and transcripts was generated. A cell-calling algorithm was used to filter the raw UMI count matrix and get the cell only filtered_feature_bc_matrix. The algorithm was similar to that of Cell Ranger and EmptyDrops^54^, which had two key steps: 1) It used a cutoff based on total UMI counts of each barcode to identify cells. This step identified the primary mode of the high RNA content cells. 2) Then the algorithm used the RNA profile of each remaining barcode to determine if it is an “empty” or a cell containing partition. This step captured the low RNA content cells whose total UMI counts might be similar to the empty wells.

### Single-cell RNA sequencing data analysis

Further quality control was applied to cells based on the following thresholds: 1) a count of expressed genes exceeding 150 but not surpassing 6,000; 2) cells containing mitochondrial RNA content lower than 10%. The DoubletFinder^55^ R package was used to remove potential doublets. The gene expression data was then processed by normalizing and scaling each sample’s filtered gene expression matrix using the functions, “NormalizeData” and “ScaleData” in the Seurat package. Batch effects across this case and other tissues were harmonized and the gene expression matrices from all samples were integrated using the Harmony^56^ R package. We performed principal component analysis (PCA) on the corrected expression matrix using highly variable genes (HVGs) identified by the “FindVariableGenes” function. The most representative principal components were used to determine different cell types with the “FindCluster” function.

### Trajectory analysis

Trajectory analysis was performed utilizing Monocle2 and Monocle3. Additionally, we employed the Python package PAGA^57^ to validate the pseudotime trajectories among 9 melanoma CTC subtypes.

### RNA-seq and data analysis

Total RNA from this case was isolated with TRIzol reagent (Invitrogen). RNA samples were subjected to RNA-seq (Novegene, Illumina Novaseq6000). The raw data were cleaned with fastp and then mapped to the hg38 genome using STAR^51^, with the following parameters: --twopassMode Basic --outReadsUnmapped None –chimSegmentMin 12 --chimJunctionOverhangMin 12 --alignSJDBoverhangMin 10 --alignMatesGapMax 200000 --alignIntronMax 200000 –chimSegmentReadGapMax parameter 3 --alignSJstitchMismatchNmax 5 -1 5 5 --runThreadN 20 --outSAMtype BAM SortedByCoordinate. The gene expression data were assembled and quantified using the RSEM package with the parameters, -paired-end and -star, and differentially expressed genes (DEGs; log2(fold change) ≥ 0.58 and P ≤ 0.05) were identified by the DEseq2^58^ R package.

### Single cell gene set enrichment analysis

We conducted the gene set enrichment analysis for select cell subtypes by the irGSEA R package, with the Hallmark or Kyoto Encyclopedia of Genes and Genomes (KEGG) pathways being derived. Finally, the “irGSEA.heatmap” and “irGSEA.halfvlnplot” functions were applied to visualize enrichment score.

### Single-cell flux estimation and cell metabolite prediction

We used scfea^59^ tools which utilizes a graph neural network model to estimate cell-wise metabolic flux by using scRNA-seq data. We chose “module gene m168” as moduleGene file, “cmMat c70 m168” as stoichiometry file, and parameter “sc imputation = True”.

### Functional and pathway enrichment analysis

Functional and pathway difference between the senescent and wild-type CTCs were performed by enrichGO with differential expression genes in two groups, enrichGO was used with log2FC >0.3 and *p* < 0.05 using the clusterProfiler R package^60^.

### Public datasets used in this study

Public available bulk RNA-seq of melanoma tissues from GSE650904, GSE19234, and GSE15605 from the Gene Expression Omnibus database (GEO, https://www.ncbi.nlm.nih.gov/geo/). Bulk RNA-seq of HES1-KD and HES1-OE MEL-167 CTCs from HRA010773 (publicly accessible at https://ngdc.cncb.ac.cn/gsa-human). Cut-tag sequence data of HES1 in MEL-167 CTCs from HRA010784. Public available scRNA-seq of 20 recurrent melanoma tissues from GSE229908. TCGA-melanoma cohort were downloaded from the UCSC Xena data portal (https://xenabrowser.net).

### Statistical analysis

Comparisons between two groups were performed using two-tailed Student’s t-test under the normality assumption. Spearman’s correlation was used to measure the correlation between two continuous variables and r > 0.3 and *P* < 0.05 was considered significant. Log-rank test was used for univariate survival analyses and showed as the Kaplan-Meier plot. All statistical analyses and visualization were performed using R or GraphPad Prism. The lines in the middle of the box plot are median and the upper and lower lines indicate 25th and 75th percentiles. *P* < 0.05 was considered statistically significant. The number of replicates and statistical tests used in figures were shown in corresponding figure legends.

## DATA AND CODE AVAILABILITY

The data generated in this study have been deposited in the Genome Sequence Archive in BIG Data Center, Beijing Institute of Genomics, Chinese Academy of Sciences, under BioProject PRJCA064084 (publicly accessible at https://ngdc.cncb.ac.cn/gsa-human). All data requests will be granted by the Data Access Committee. scRNA-seq are also available in GSA with the accession numbers HRA018444. Bulk RNA-seq are also available in GSA with the accession numbers HRA010772 and HRA010773.Cut-tag sequencing data are also available in GSA with the accession numbers HRA010784. The detailed information of all public datasets has been provided in **Supplementary Table S10**.

No original code was generated in this study. Statistical analyses were performed using standard R packages as described below.

Any additional information required to reanalyze the data reported in this article is available upon request.

## ACKNOWLEDGMENTS

We are grateful to all the patients who participated by donating specimens for this study. This study was supported by the National Key R&D Program of China: No. 2023YFC2705804 (to X.H), National Natural Science Foundation of China (No. 82573274 to X.H, No. 82422062 and 82273182 to WN. G), Guangdong provincial funding awards: 2021QN02Y112, 2023A1515010287 (to X.H). Translational research funding from SUSTech SOM-ShangYiJian Joint Laboratory:20230004056 (to X.H). Guangdong Basic and Applied Basic Research Foundation: NO. 2022A1515140187 (to M.X).

## AUTHOR CONTRIBUTIONS

**G. Huang**: Conceptualization, data curation, software, visualization, formal analysis, investigation, methodology, writing–original draft, writing–review and editing. **X. Xu**: Data curation, formal analysis, investigation, methodology, writing–original draft, writing–review and editing. **B. Zhang**: Data curation, formal analysis, investigation, methodology, writing–original draft, writing–review and editing. **M. Zhao**: Formal analysis, resources, visualization. **Y. Cheng**: mIHC experimental analysis. **B. Zhao**: Cell culture methodology and analysis. **S. Zheng, S. Yu**: Visualization, methodology and analysis. **W. Liu, J. Hu:** Formal analysis, investigation, methodology. **C. Long:** Reporter construction, methodology and data analysis. **X. Liu**: Software, methodology and data analysis. **Y. Zhang**, **Y. Sheng**, **S. Xia, J. Liu**: Methodology and data analysis. **L. Zeng, H. Yu, H. Yang**: microfluidic CTC isolation technology support and guidance. **Y. Lu, J. Zhang, W. Feng, M. Xu, W. Guo**: Resources, supervision, project administration, writing–review and editing. **X. Hong**: Conceptualization, supervision, data analysis, funding acquisition, writing–original draft, project administration, writing–review and editing.

---

**Supplementary Figure S1.**
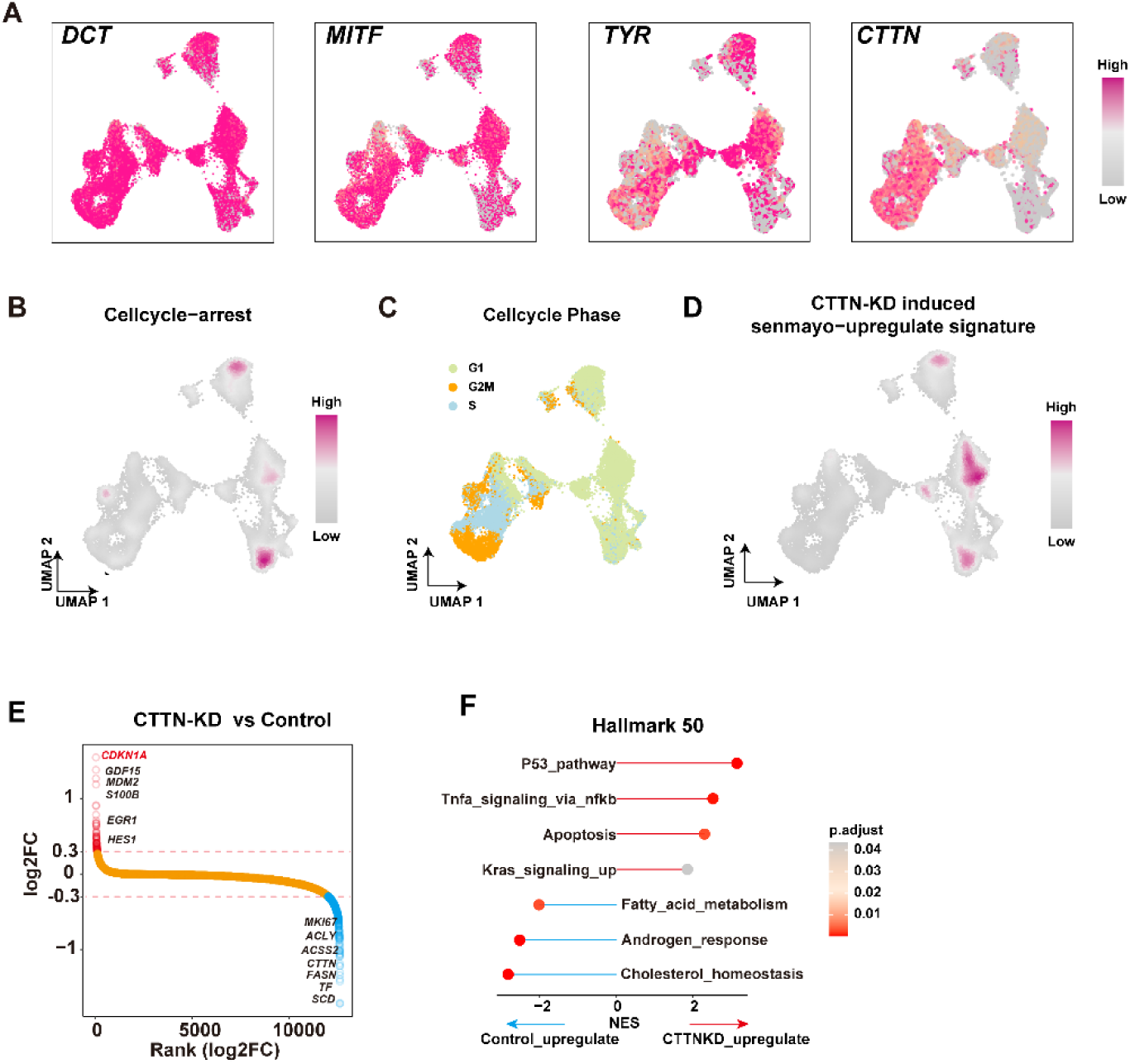
Differentially expression genes and pathways in CTTN-KD induced senescent CTCs. (A) UMAP shows mRNA expression levels of melanocyte lineage associated markers (DCT, MITF, TYR) and CTTN among wild-type and senescent CTCs subpopulations. (B-D) UMAP shows the pathways (Cellular cycle arrest (B), Cell cycle (C) and CTTN-KD induced Senmayo upregulate signatures (D)) among wild-type and senescent CTCs subpopulations. (E) Differentially expression genes among wild-type and senescent CTCs subpopulations. The *y*-axis indicates the log2FC (fold change) and genes ordered by log2FC along the x-axis. (F) Pathway enrichment of wild-type and senescent CTC subpopulations. NES, normalized enrichment score.

**Supplementary Figure S2.**
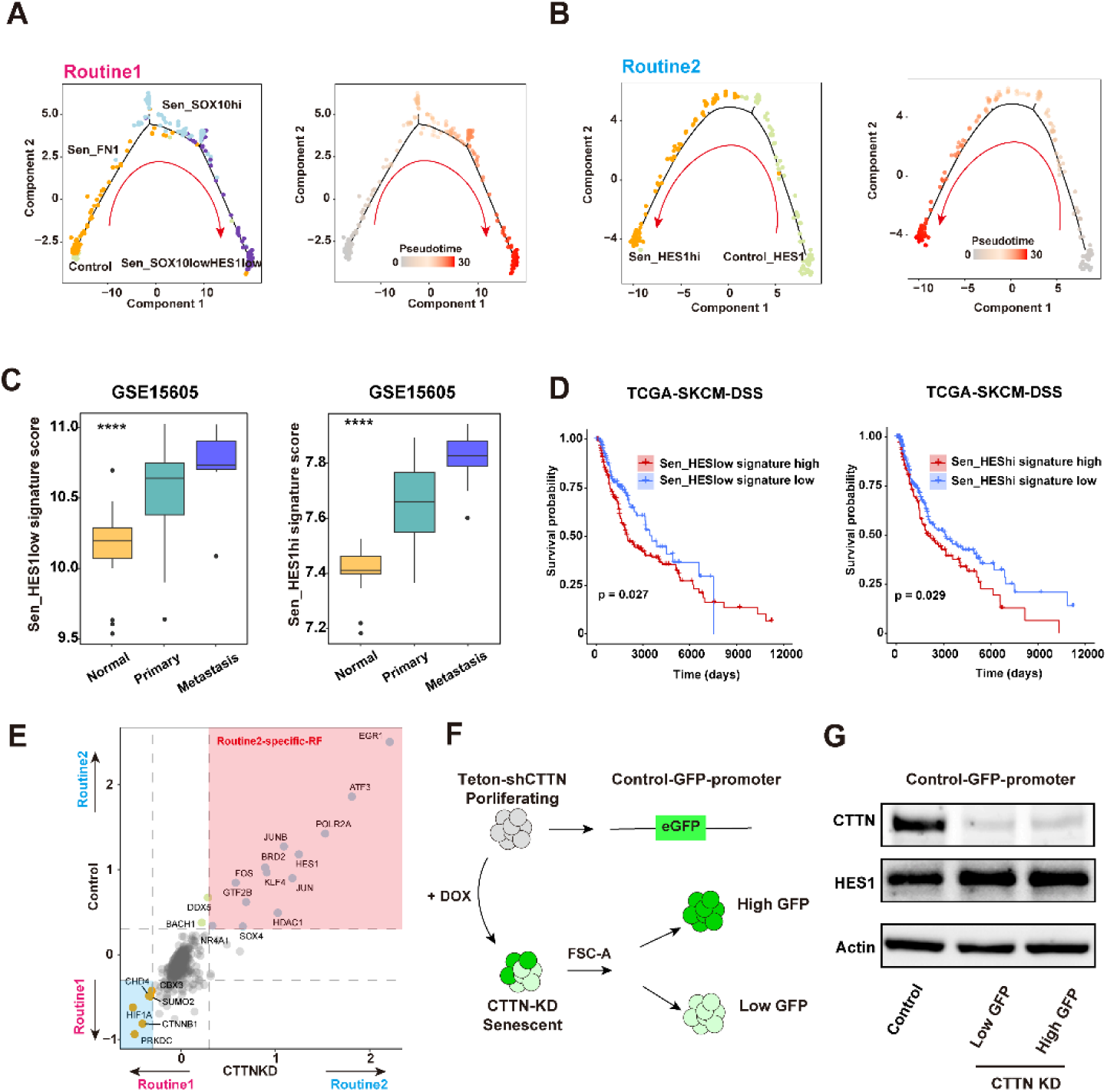
Trajectory analysis and clinical relevance of senescent CTC subpopulations. (A) Trajectory analysis of wild-type and senescent melanoma CTCs in route1 was performed by Monocle 2, colored by inferred pseudotime. (B) Trajectory analysis of wild-type and senescent melanoma CTCs in routine2 was performed by Monocle 2, colored by inferred pseudotime. (C) Boxplot shows the expression level of Sen_HES1^low^ and Sen_HES1^high^ subpopulation signatures in primary tumor, metastasis, and normal tissue of melanoma patients in the GSE15605 cohort. (D) Disease specific survival (DSS) of melanoma patients in The Cancer Genome Atlas Program (TCGA) based on Sen_HES1^low^ and Sen_HES1^high^ subpopulations signatures expression. (E) Quadrant analysis shows the overlap of differentially expressed regulator factors in route1 versus route2 in wild-type and senescent melanoma CTCs. (F) Schematic diagram of Control-eGFP reporter system to isolate the GFP^high^ and GFP^low^ senescent CTCs following CTTN-KD in CTCs. (G) Protein levels of CTTN and HES1 in GFP^high^ and GFP^low^ senescent CTCs and wild-type CTCs. Actin as a loading control. **C,** Data are represented as mean ± SEM. **C,** statistical significance was calculated by Kruskal-Wallis test. **D,** statistical significance was calculated by log-rank test. NS, not significant; *, *P* < 0.05; ****, *P* < 0.0001.

**Supplementary Figure S3.**
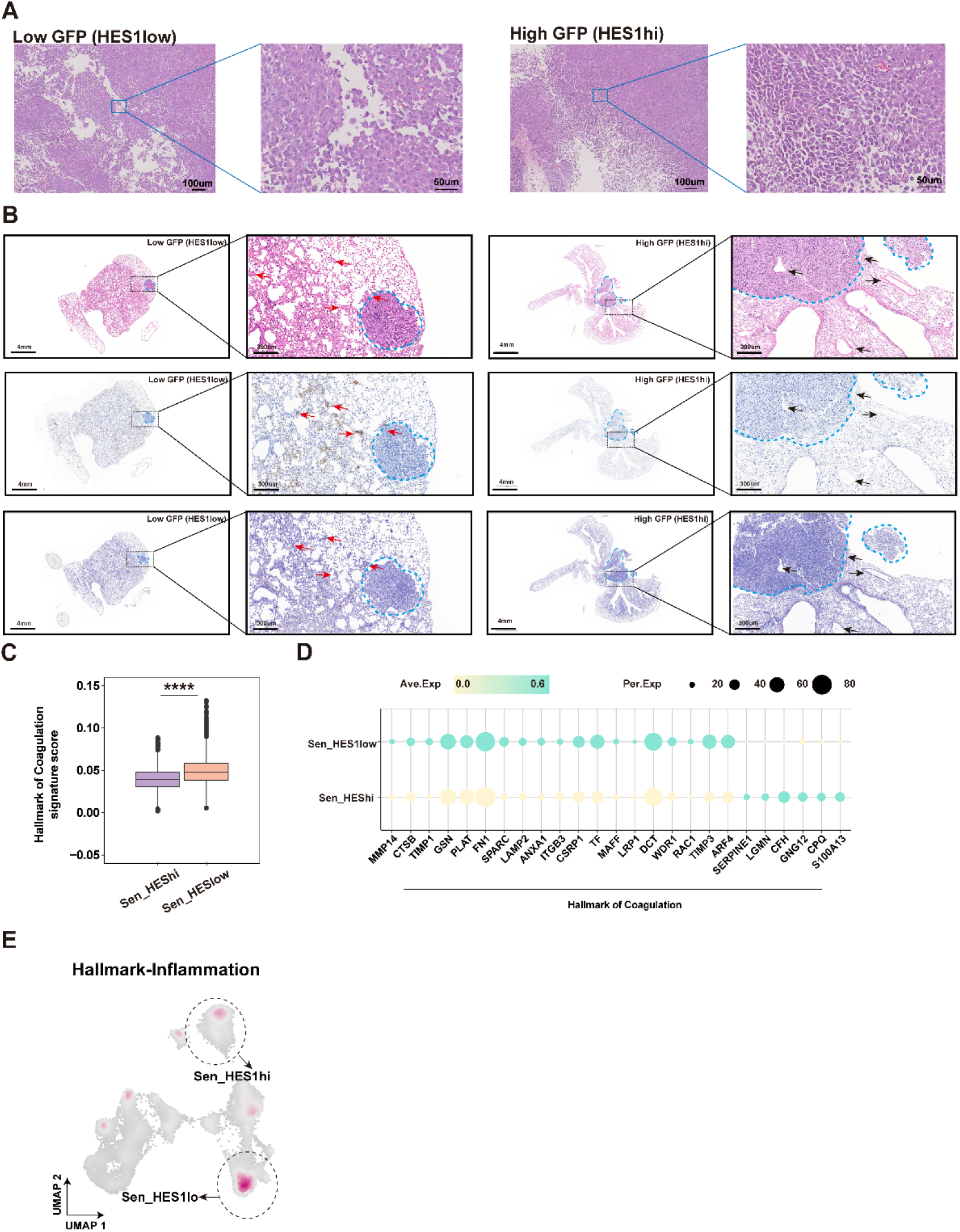
Metastatic relapse heterogeneity of senescent melanoma CTCs. (A) Representative images of H&E staining of HES1^low^ and HES1^high^ relapse tumors in **Figure 3E**. (B) Representative images of H&E staining (top), IHC staining (middle) and PATH (phosphotungstic acid-haematoxylin) staining (bottom) in lung tissues of HES1^low^, HES1^high^ groups in **Figure 3F**. (C) Bar graph shows coagulation pathway signature scores in Sen_HES1^low^ and Sen_HES1^high^ subpopulations. (D) Dot plot shows coagulation associated genes expressed in Sen_HES1^low^ and Sen_HES1^high^ subpopulations. Dot size indicates the fraction of expressing cells, and the colors represent normalized gene expression levels. (E) UMAP shows inflammation pathway among wild-type and senescent CTC subpopulations. **C,** Data are represented as mean ± SEM. **C,** statistical significance was calculated by Wilcoxon test. ****, *P* < 0.0001.

**Supplementary Figure S4.**
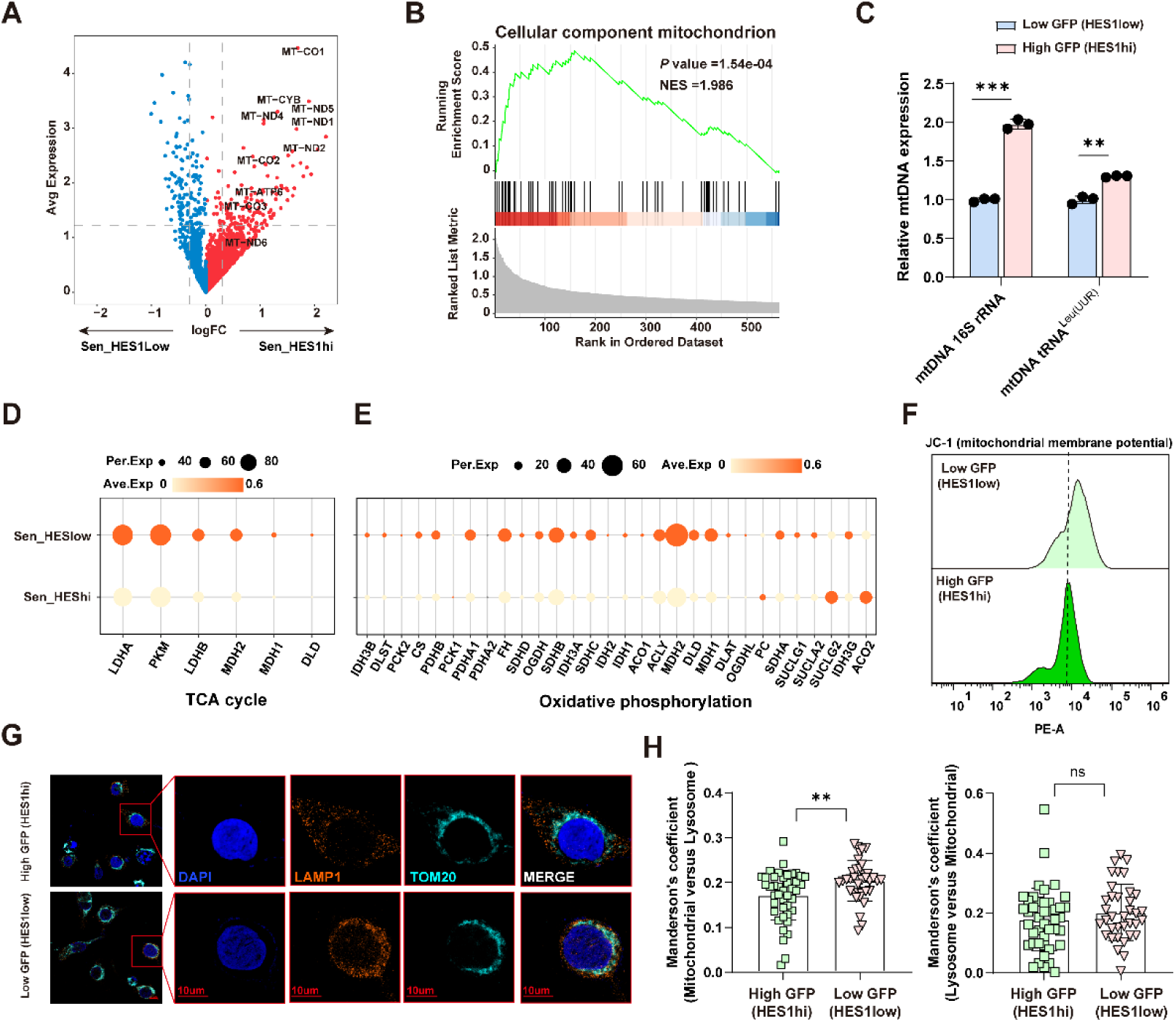
Molecular feature diversity within senescent CTCs. (A) Volcano plot of differentially expression genes of Sen_HES1^low^ and Sen_HES1^high^ subpopulations. (B) GSEA shows the GO cellular component of mitochondrial enriched in Sen_HES1^high^ subpopulations. (C) Relative mtDNA copy number determined by qPCR in HES1^low^, HES1^high^ senescent CTCs. *n* =3 independent experiments. (D) Dot plot shows TCA cycle associated genes expressed in Sen_HES1^low^ and Sen_HES1^high^ subpopulations. Dot size indicates the fraction of expressing cells, and the colors represent normalized gene expression levels. (E) Dot plot shows oxidative phosphorylation associated genes expressed in Sen_HES1^low^ and Sen_HES1^high^ subpopulations. Dot size indicates the fraction of expressing cells, and the colors represent normalized gene expression levels. (F) Flow cytometric analysis of JC-1 staining shows mitochondrial membrane potential in HES1^low^ and HES1^high^ cells. (G-H) Representative images and quantification analysis of colocalization of LAMP1 and TOM20 in HES1^low^, and HES1^high^ senescent CTCs. Manders’s coefficient (mitochondria versus lysosomes) indicates the proportion of mitochondria involved interaction with lysosome in total mitochondria. Manders’s coefficient (Lysosome versus Mitochondria) indicates the proportion of lysosomes involved interaction with mitochondria in total lysosomes. **C,** and **H,** Data are represented as mean ± SEM. **C,** and **H** Statistical significance was calculated by a two-sided Student *t* test; NS, not significant; **, *P* < 0.01; ***, *P* < 0.001. mtDNA, mitochondrial encoded DNA.

**Supplementary Figure S5.**
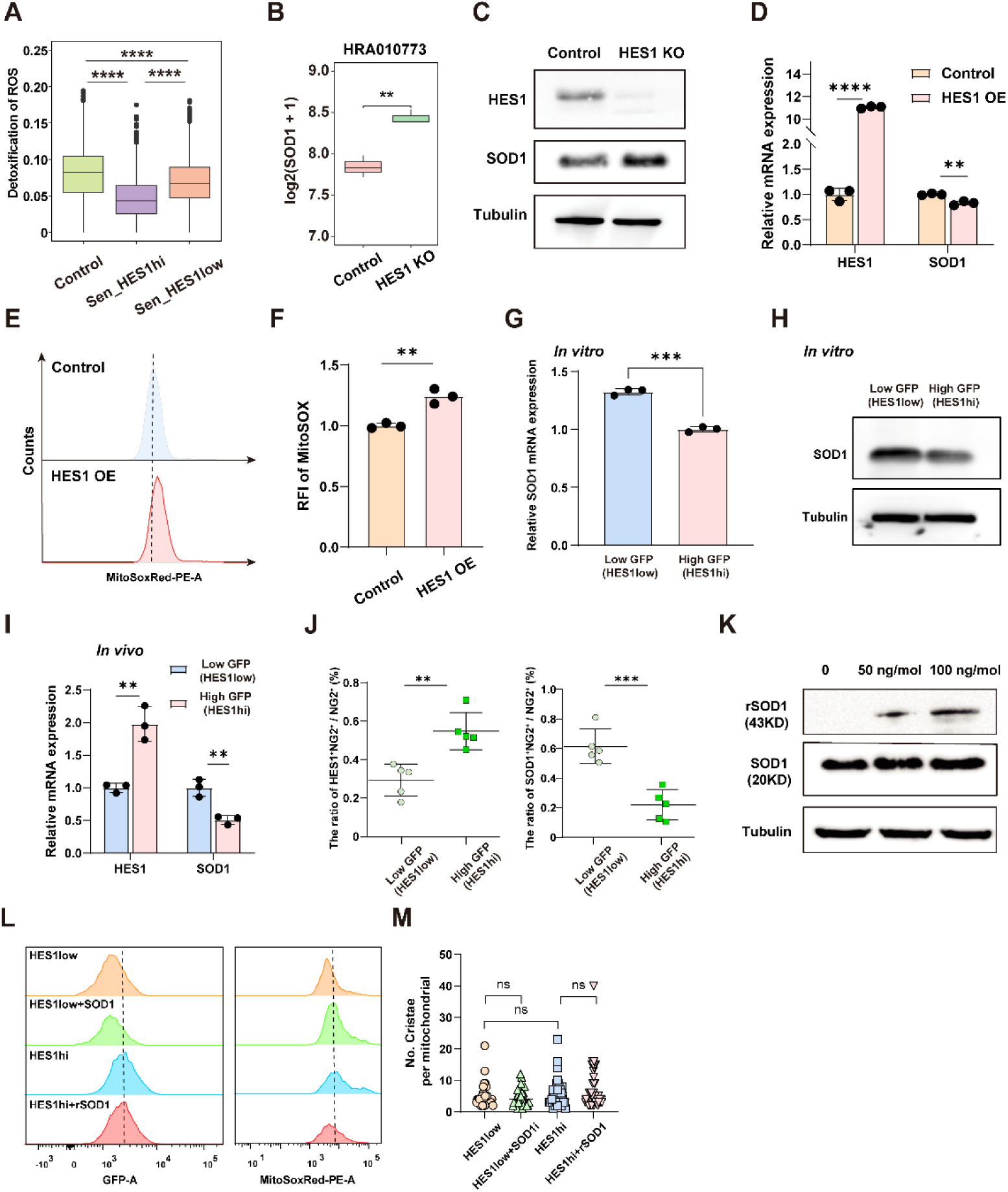
HES1 negatively regulates SOD1 in melanoma Mel-167 CTCs. (A) Box plot shows the detoxification of ROS pathway in control, Sen_HES1^low^, and Sen_HES1^high^. (B) Box plot shows the mRNA expression of SOD1 in *HES1*-knockout CTCs and wild-type CTCs from HRA010773 dataset. (C) Protein levels of HES1 and SOD1 in *HES1*-knockout CTCs and wild-type CTCs. Tubulin as a loading control. (D) Relative mRNA expression levels of *HES1* and *SOD1* determined by RT-qPCR in *HES1*-knockout CTCs and wild-type CTCs. *n* = 3 independent experiments. (E) Flow cytometry analysis of Mitosox Red in *HES1*-knockout CTCs and wild-type CTCs. (F) Quantification analysis of Mitosox Red in *HES1*-knockout CTCs and wild-type CTCs. *n* = 3 independent experiments. (G) Relative mRNA expression of *SOD1* determined by RT-qPCR in HES1^low^, HES1^high^ senescent CTCs. *n* = 3 independent experiments. (H) Western blotting shows protein levels of SOD1 in HES1^low^ and HES1^high^ senescent CTCs. Tubulin as loading control. (I) Relative mRNA expression of *HES1* and *SOD1* determined by RT-qPCR in HES1^low^, HES1^high^ senescent tumors in **Figure 3E**. *n* = 3 independent experiments. (J) Quantification analysis of the proportion of HES1^+^NG2^+^ and SOD1^+^NG2^+^ cells in NG2^+^ cells in HES1^low^, HES1^high^ tumors from **Figure 3E**. *n* = 5 independent experiments. (K) Protein levels of SOD1-GST and SOD1 in HES1^low^ senescent CTCs with or without SOD1-GST treatment. Tubulin as loading control. (L) Flow cytometry analysis of GFP and MitoSOX Red in HES1^low^ senescent CTCs, HES1^high^ senescent CTCs, HES1^low^ senescent CTCs with LCS-1(SOD1 inhibitor) treatment, and HES1^high^ senescent CTCs with SOD1 recombinant protein treatment. *n* = 3 independent experiments. (M) Quantification of the number of cristae per mitochondrion of HES1^low^ senescent CTCs, HES1^high^ senescent CTCs, HES1^low^ senescent CTCs with LCS-1(SOD1 inhibitor) treatment, and HES1^high^ senescent CTCs with SOD1 recombinant protein treatment. **A, B, D, F, G, I, J,** and **M,** Data are represented as mean ± SEM. **A, B, D, F, G, I, J,** and **M,** Statistical significance was calculated by a two-sided Student *t* test; **, *P* < 0.01; ***, *P* < 0.001; ****, *P* < 0.0001. rSOD1(SOD1-GST), GST tagged SOD1 recombinant protein. PE, phycoerythrin

**Supplementary Figure S6.**
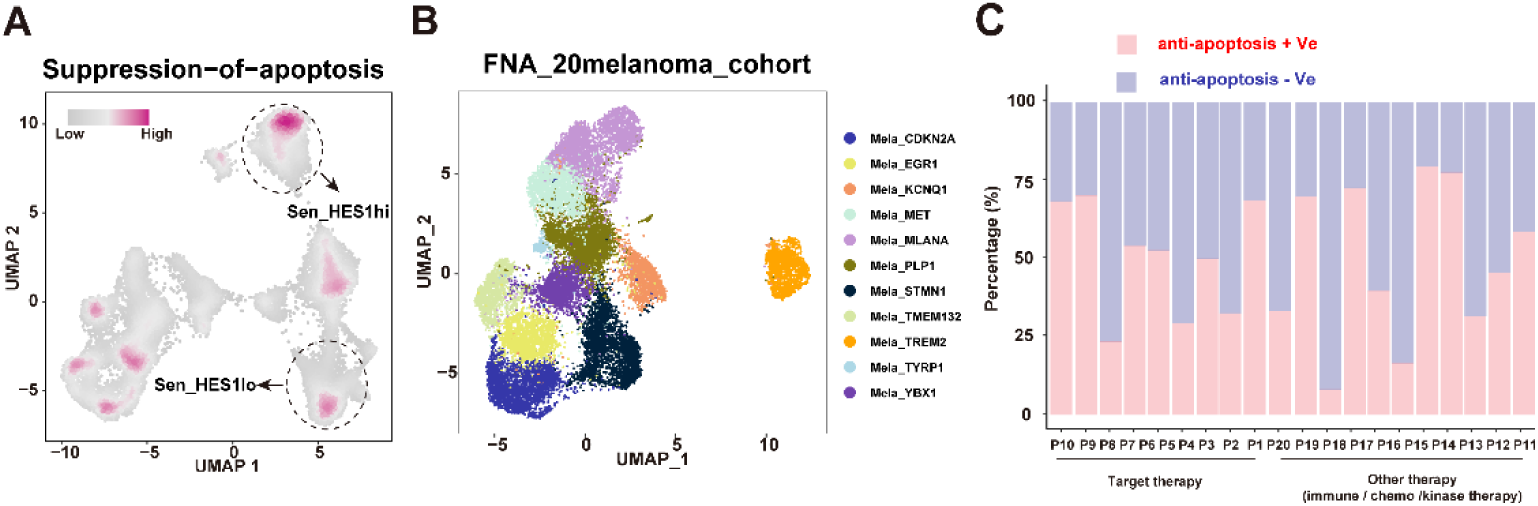
BCL-2 family signatures are enriched at relapse and suggest potential therapeutic vulnerability. (A) UMAP shows the suppression of apoptosis pathway among wild-type and senescent CTC subpopulations. (B) UMAP visualization of subpopulations of relapsed melanoma in 20 patients. Each dot indicates a single cell. Color-coded for clusters. (C) Bar plot shows relative proportions of anti-apoptosis signatures positive (anti-apoptosis + Ve) and anti-apoptosis signatures negative (anti-apoptosis - Ve) melanoma cells in each patient.

**Supplementary Figure S7.**
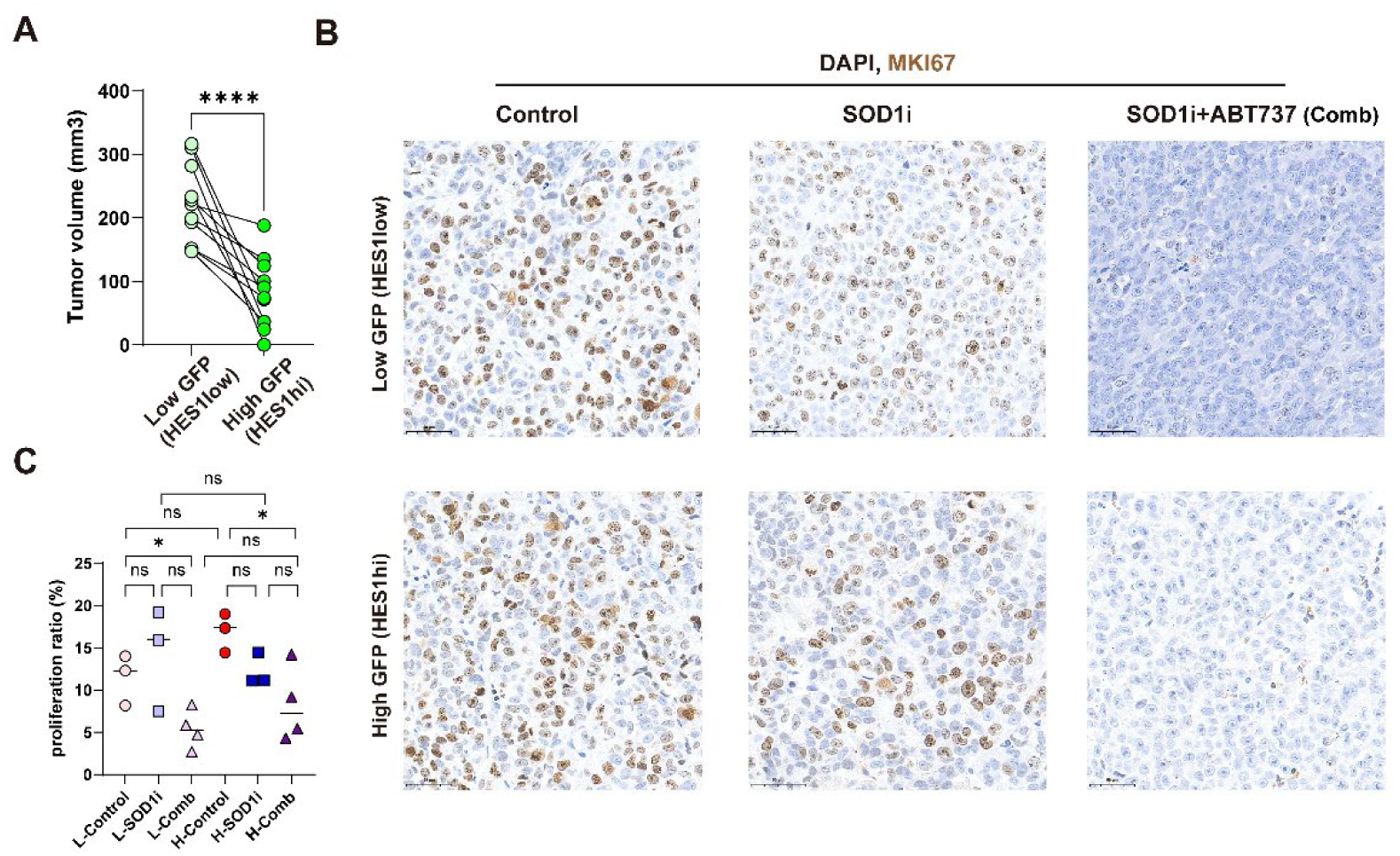
Combination therapy with SOD1 inhibitor and senolytic drug effectively inhibits proliferative potential of HES1^low^ and HES1^high^ relapse tumors *in vivo*. (A) Quantification analysis of tumor volume of HES1^low^ and HES1^high^ relapse tumors at day0. (B) Representative images of proliferating cells via MKI67 staining of HES1^low^ and HES1^high^ relapse tumors in Control, SOD1i and Comb groups. Magnification, x40. L-Control, L-SOD1i, and L-Comb refer to HES1^low^ relapse tumors in Control, SOD1i and Comb groups, respectively. H-Control, H-SOD1i, and H-Comb refer to HES1^high^ relapse tumors in Control, SOD1i and Comb groups, respectively. (C) Quantification of proliferating cells via MKI67 staining of relapse tumors from **Figure S7B. A,** and **C,** Data are represented as mean ± SEM. **A,** Statistical significance was calculated by two-tailed Student’s t-test. **C,** Statistical significance was calculated by two-way ANOVA with the Bonferroni *post hoc* test. NS, not significant; *, *P* < 0.05; ****, *P* < 0.0001.

**Supplementary Figure S8.**
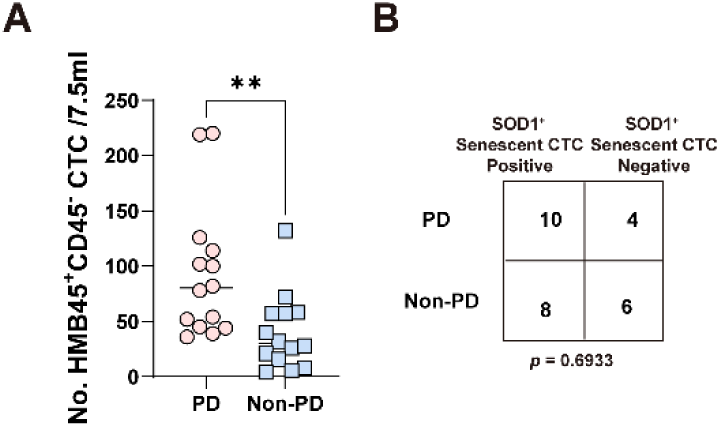
CTCs are associated with clinical status of first-line, on-treatment melanoma patients. (A) Box plot shows the number of HMB45^+^ CD45^-^ CTCs in 7.5mL blood within PD and non-PD melanoma patients. (PD, progressive disease, *n* =14; Non-PD, stable disease or partial response, *n* =14). (B) Fisher test was performed to assess the association between β-gal^+^SOD1^+^ senescent CTCs and clinical outcome. β-gal^+^HMB45^+^CD45^-^ SOD1^+^ cells were recognized as SOD1^+^ senescent CTCs. **A,** and **B,** Data are represented as mean ± SEM. **A,** Statistical significance was calculated by a two-sided Student *t* test; **B,** statistical significance was calculated by Fisher test. **, *P* < 0.01.

